# Cloning and functional characterization of novel human neutralizing anti-interferon-alpha and anti-interferon-beta antibodies

**DOI:** 10.1101/2024.05.05.591636

**Authors:** Emmanouil Papasavvas, Lily Lu, Matthew Fair, Isabela Oliva, Joel Cassel, Sonali Majumdar, Karam Mounzer, Jay R. Kostman, Pablo Tebas, Amit Bar-Or, Kar Muthumani, Luis J. Montaner

## Abstract

Type I interferons (IFNs) play a pivotal role in immune response modulation, yet dysregulation is implicated in various disorders. Therefore, it is crucial to develop tools that facilitate the understanding of their mechanism of action and enable the development of more effective anti-IFN therapeutic strategies. In this study, we isolated, cloned, and characterized anti-IFN-α and anti-IFN-β antibodies (Abs) from peripheral blood mononuclear cells of individuals treated with IFN-α or IFN-β, harboring confirmed neutralizing Abs. Clones AH07856 and AH07857 were identified as neutralizing anti-IFN-α-specific with inhibition against IFN-α2a, -α2b, and -αK subtypes. Clones AH07859 and AH07866 were identified as neutralizing anti-IFN-β1a-specific signaling, and able to block Lipopolysaccharide or S100 calcium binding protein A14-induced IFN-β signaling effects. Cloned Abs bind rhesus but not murine IFNs. The specificity of inhibition between IFN-α and IFN-β suggests potential for diverse research and clinical applications.

## INTRODUCTION

Interferons (IFNs) are a family of widely expressed and related cytokines that have potent antiviral, antiproliferative, antitumor and immunomodulatory activities ^1–4^. They act by inducing innate and promoting adaptive immune responses, and thus serve as the first line of defense against viral, bacterial infections, and malignant cells ^5,6^. The IFN family consists of distinct proteins that fall into three discrete classes (type I, II and III IFNs) differentiated by their receptor complexes ^7–11^.

The type I IFN family includes 13-14 IFN-α subtypes which are involved in the innate immune response to infection, tumor development, and other inflammatory stimuli ^12,13^. All the members of the type I IFN family bind to a common heterodimeric receptor (IFNα/β receptor, IFNAR) consisting of the low affinity IFNAR1 and the high affinity IFNAR2 subunits ^6,14,15^. Binding of type I IFNs to IFNAR2 results in the subsequent recruitment of IFNAR1, which leads to the formation of a high-affinity IFNAR. Although all type I IFN subtypes start signaling through the same type I IFNAR, the stability of binding between various type I IFN subtypes with IFNAR is different. IFN-α1 binds with the lowest affinity to both chains of IFNAR, while IFN-α2 binds to IFNAR1 with lesser (micromolar) affinity than to IFNAR2 (nanomolar) ^16^. On the other hand, IFN-β has a greater affinity for IFNAR1 and can also bind to IFNAR1 independently of IFNAR2 leading to transduction of specific, unconventional intracellular signals and contributing to toxicity in vivo ^6,17^.

Binding of type I IFNs to IFNAR results in the subsequent activation mainly of the Janus kinase (JAK)-signal transducer and activator of transcription (STAT) (JAK-STAT) pathway, and the transcription of IFN-stimulated genes [ISG, e.g. MX Dynamin Like GTPase 1 (MX1), Ubiquitin specific peptidase 18 (USP18)], which in turn allows cells to respond to IFN stimulation ^4,18–22^. The activities of type I IFNs can be grouped to robust, and tunable. Robust activities (e.g. activities associated with anti-viral responsiveness) are common to all cells, require only minute amounts of all type I IFNs, and are independent of their binding affinity or the numbers of IFNAR on cell surface. On the other hand, tunable activities (e.g. activities observed after activation of robust activities and associated with the immunomodulatory and anti-proliferative effects of type I IFNs) are cell type-specific, require 1000-fold higher concentration of type I IFNs and higher number of IFNAR, are observed after longer times of treatment with type I IFNs, and are most strongly activated by the high affinity binding of IFN-β (or IFN-α2 variants engineered for tight binding) to IFNAR ^15^.

Type I IFNs have been used widely in the clinic primarily in the treatment of viral infections and of autoimmune conditions such as multiple sclerosis (MS). Regarding viral infections, IFN-α2a/b has been used to treat hepatitis virus B and C ^23–25^. In addition, in the context of HIV infection treatment with IFN-α2a/b was high in the pre-antiretroviral therapy (ART) era ^26,27^ but was abandoned after wide usage of ART. Recent reports though, suggest a potential beneficial role of IFN-α2a/b in activating natural killer cell (NK) anti-viral activity in HIV cure-directed strategies ^28–32^. While antiviral strategies have pursued robust activities of type I IFNs via IFN-α2a/b, therapy for MS has focused on tunable activities of decreasing T cell activation as exemplified by usage of IFN-β ^33,34^. Therapy with type I IFNs can result in the development of natural anti-IFNs antibodies (Abs) such as natural anti-IFN-β Abs in patients treated with IFN-β for MS that block IFN-β but do not block IFN-α ^35–40^, as well as development of natural anti-IFN-α Abs in patients treated with IFN-α. ^41–48^

Despite the central role of type I IFNs in the development of immune response and their great clinical importance, aberrant activation of the type-I IFN response has been shown to result in a large spectrum of disorders called interferonopathies, while in the context of HIV and SIV infection type I IFNs have been described to have beneficial and detrimental roles ^5,26,49–56^. Limited information has defined whether these differential outcomes are linked to specific type I IFN subtypes. However, in both viral and cancer a potential detrimental role has been linked to IFN-α or IFN-β expression ^53,57–60^ supporting development of novel strategies for the selective inhibition of specific type I IFNs signaling. While human studies targeting in the same study specific type I IFN subtypes IFN-α versus IFN-β have yet to be performed, a larger number of studies has been conducted testing the effectiveness of inhibitors of all type-I IFN signal transduction, of monoclonal Abs (mAbs) against IFN-α, and of IFNAR antagonists ^61–65^. Here, we identify, clone and characterize novel natural neutralizing human anti-IFN-α and anti-IFN-β Ab clones derived from a single pooled peripheral blood mononuclear cells cDNA from persons treated with IFN-α or IFN-β after having confirmed the respective presence of plasma neutralizing anti-IFN-specific Abs.

## RESULTS

### Identification of neutralizing Abs specific against IFN-β or IFN-α

We used the IFNR dimerization assay in the presence of IFN-β or IFN-α respectively, as a primary screen for identification of persons carrying neutralizing IFN-specific plasma Abs. First, MS patient plasma with a history of treatment with IFN-β were analyzed for anti-IFN-β activity. We identified two plasmas with ability to inhibit IFN-β-mediated activity (MS-001, and MS-006) in contrast to other MS plasma or non-MS control plasma M486-B55 or F623-C32 (**Figure 1A**). Due to sample availability, we further tested MS-001 plasma inhibitory effect in the IFN-β dimerization assay using different concentrations of IFN-β (**Figure 1B top panel**) or different concentrations of plasma and IFN-β (**Figure 1B bottom panel**). MS-001 plasma showed no inhibition in the IFN-α dimerization assay even when lower concentrations of IFN-α were used together with the highest plasma concentration (11% final in the well) (**Figure 1C**). To confirm that the observed activity was associated with anti-IFN-β IgG in plasma, sequential IgG depletions from MS-001 plasma were completed resulting in the loss of inhibitory effect in the IFN-β dimerization assay (**Figure 1D**). It should be noted that as shown in **Figure S1**, the depletion of Abs was specific as no depletion of albumin in the same plasma samples was observed.

**Figure 1.**
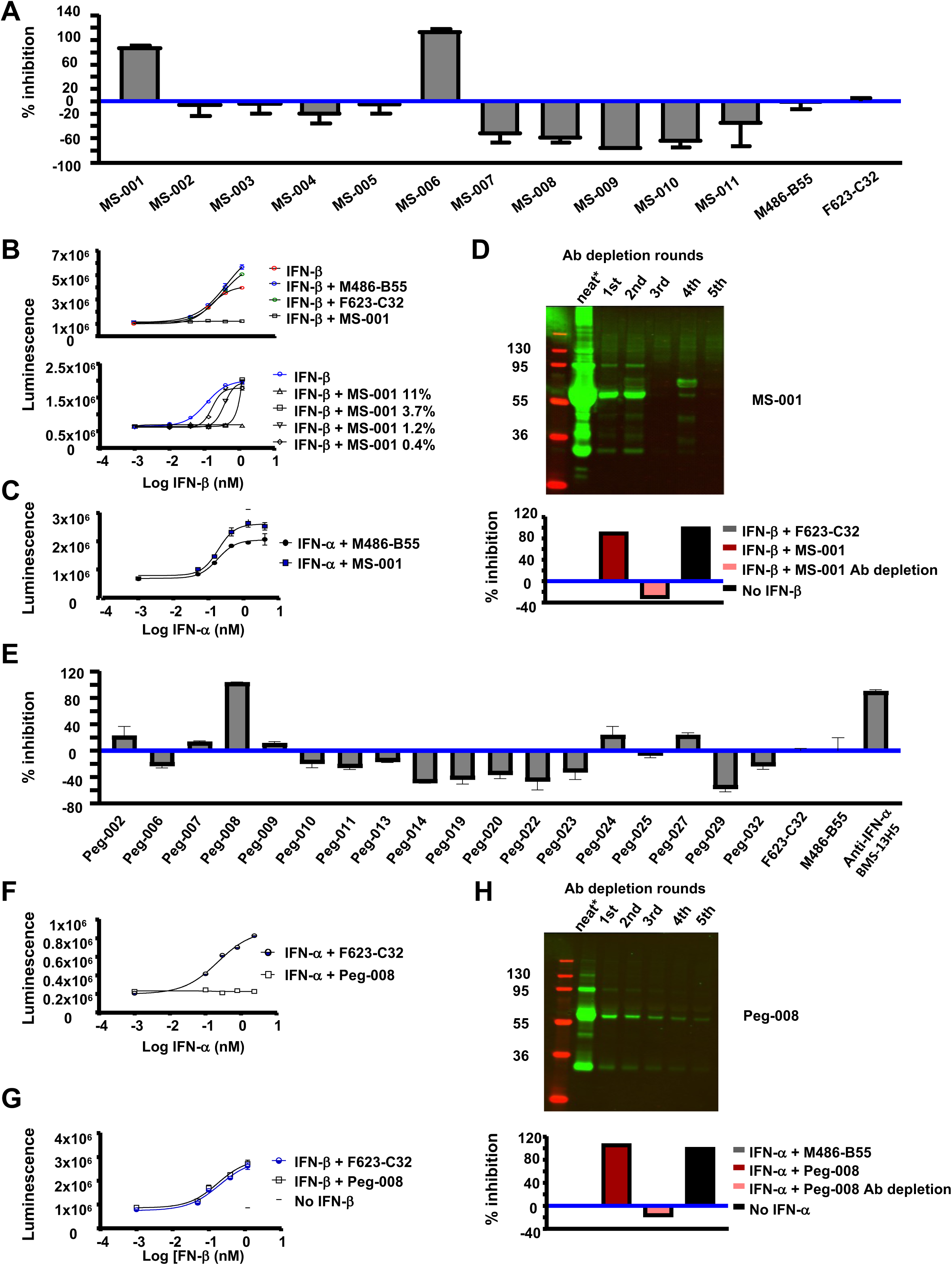
Identification of neutralizing IFN-α-specific and IFN-β-specific plasma from IFN-α or IFN-β-treated persons. (A) Percent (%) inhibition of IFNAR dimerization in the presence of IFN-β and plasma from participants with multiple sclerosis (MS, MS-001 to MS-011) or from healthy control individuals M486-B55, F623-C32. Plasma was pre-incubated overnight with IFN-β at 4°C, at 3X of the final concentrations (final concentrations in the wells with HEK IFNAR1/IFNAR2 cells: 11% plasma, 1.2 nM IFN-β). (B) Top panel shows luminescence for IFNAR dimerization by IFN-β (in limiting dilutions) in the presence or absence of 11% plasma from participant MS-001 or from healthy control individuals M486-B55 and F623-C32. Bottom panel shows luminescence for IFNAR dimerization by IFN-β (in limiting dilutions) in the presence or absence of plasma (in limiting dilutions) from participant MS-001. (C) Luminescence for IFNAR dimerization by IFN-α (in limiting dilutions) in the presence or absence of 11% plasma from participant MS-001 or from healthy control individual M486-B55. (D) Top panel shows western blot of 10% plasma from participant MS-001 following 5 rounds of Ab depletion with A/G sepharose beads. Bottom panel shows % inhibition of IFNAR dimerization in the absence of 1.2nM IFN-β, or in the presence of 1.2nM IFN-β and 10% plasma from participant MS-001 (before and after Ab depletion) or from healthy control individual F623-C32. See Figure S1 for specificity of Ab depletion in plasma from participant MS-001. (E) Percent (%) inhibition of IFNAR dimerization in the presence of IFN-α and plasma from ART-suppressed persons living with HIV having a history of receiving IFN-α immunotherapy (Peg-002 to Peg-032), or plasma from healthy control individuals M486-B55, F623-C32, or neutralizing anti-IFN-α Ab (BMS-13H5)]. Plasma was heat inactivated at 56°C for 1 hr and then pre-incubated overnight with IFN-α at 4°C, at 3X of the final concentrations (final concentrations in the wells with HEK IFNAR1/IFNAR2 cells: 11% plasma, 2.4 nM IFN-α, 1.1. μg/ml anti-IFN-α Ab). (F) Luminescence for IFNAR dimerization by IFN-α (in limiting dilutions) in the presence or absence of 11% plasma from participant Peg-008 or from healthy control individual F623-C32. (G) Luminescence for IFNAR dimerization by IFN-β (in limiting dilutions) in the presence or absence of 11% plasma from participant Peg-008 or from healthy control individual F623-C32. (H) Top panel shows western blot of 10% plasma from participant MS-001 following 5 rounds of depletion of Abs with A/G sepharose beads. Bottom panel shows % inhibition of IFNAR dimerization in the absence of 2.4nM IFN-α, or in the presence of 2.4nM IFN-α and 10% plasma from participant Peg-008 (before and after Ab depletion) and from healthy control individual M486-B55. In panels (B), (C), (F) and (G) plasma was heat inactivated at 56°C for 1 hr and then pre-incubated overnight with IFN-β or IFN-α at 4°C, at 3X of the final concentrations. Dilution values shown are the final concentrations in the wells with cells. Data in panels (A), (D), (E) and (H) are shown as means and STDEV of % inhibition.

For activity against IFN-α among ART-suppressed persons living with HIV, only plasma from Peg-008 showed the highest inhibitory effect in the IFN-α dimerization assay even when lower concentrations of IFN-α were tested (**Figure 1E, F**). No inhibitory effect was found against IFN-β in same assay by Peg-008 plasma (**Figure 1G**). As above, activity against IFN-α in Peg-008 plasma was lost after Ab depletion from plasma (**Figure 1H**). Overall, plasmas with IgG-mediated neutralizing anti-IFN-specific activity were identified.

### Identification and cloning of sequence of diverse IFN-α and IFN-β binding Abs using pooled donor cDNAs into Ab phage display technology

To clone the neutralizing anti-IFN-α and IFN-β Abs identified above, a workflow based on Ab phage display technology from PBMC-cell cDNA as starting material was employed (**Figure S2A**). Using PBMC, complementary DNA (cDNA) was synthesized and pooled from mRNA created joining MS-001 and Peg-008. The variable region gene segments (VH and VL) of the heavy and light chains were then amplified via polymerase chain reaction. These VH and VL gene segments were subsequently cloned into specialized phagemid expression vectors, generating a library of single-chain Fv (scFv) DNAs, where "Fv" refers to the variable region fragment of an Ab. Through 2-4 rounds of panning against either IFN-α or IFN-β, remaining scFv/phage with the ability to selectively bind to a single IFN were identified. From bacterial colonies, individual scFv/phage (referred to as monoclonal phage) were isolated, enabling the identification of the top six specific binding clones (3 against IFN-α: AH07808, AH07856, AH07857, 3 against IFN-β: AH07859, AH07866, AH07882) and 5 clones with partial binding activity (AH07824, AH07831, AH07845, AH07858, AH07881).

The VH and VL DNA sequences of selected scFv/phages were subjected to sequencing for VH and VL fragment, and subsequently subcloned into expression plasmids that incorporated Ab constant region domains (**Figure 2A)**, thereby reconstituting authentic bivalent IgG Abs. The nucleotide sequences of the top six anti-IFN-α2a and anti-IFN-β mAbs were analyzed for diversity by phylogenic trees based on the sequences of complementarity-determining regions (CDRs). Included in analysis were added partially reactive clones as well as nucleotide sequence of previously reported neutralizing Abs against IFN-α (BMS-13H5; patent W02005059106A2) or IFN-β (CTI-AF1; patent US20170313769A1) respectively ^66,67^. Notably, diversification of CDRs was observed between the newly identified clones against IFN-α (**Figure S2B**) or anti-IFN-β (**Figure S2C**) and previously reported neutralizing Abs. This study presents a novel method for the identification of Abs selectively binding to IFN-α or IFN-β. Leveraging a multiple-persons pooled cDNA approach, clones with variable regions of both kappa (VK) and lambda (VL) chains were isolated. These clones demonstrated efficacy in selectively binding to IFN-α or IFN-β mAbs. The results underscore the effectiveness of this approach in cloning diverse and specific CDRs targeting IFN-α and IFN-β. This method offers advantages in capturing a wide range of Ab specificities and can expedite the discovery and development of therapeutic Abs for various clinical applications.

**Figure 2.**
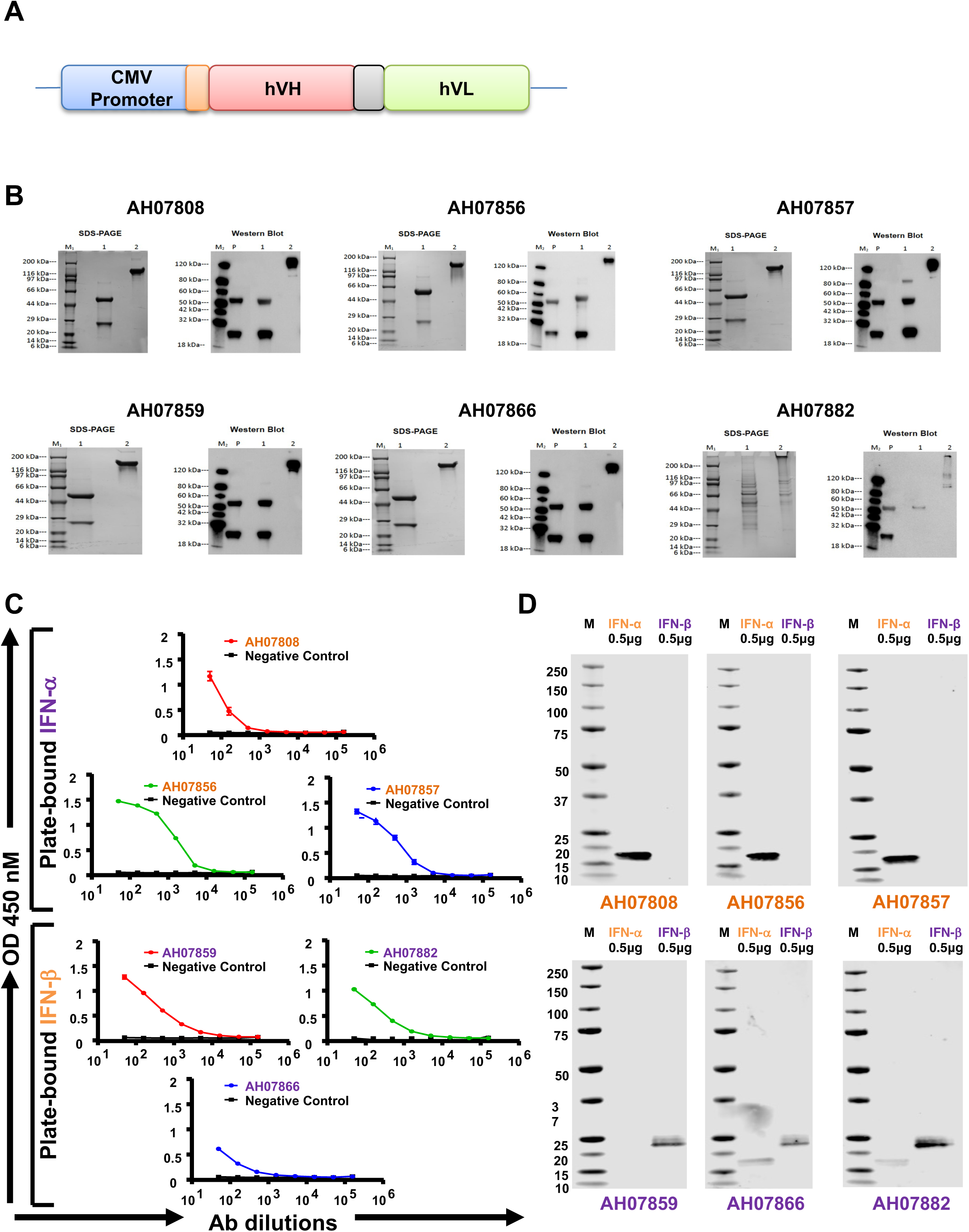
Characterization of expressed and purified mAbs using Western blot and Coomassie blue straining together with analysis of mAb reactivities with IFN-α2a and IFN-β proteins using ELISA and Western blot assays. (A) The amplified VH and VL genes were selectively assembled into DNA cassettes. The pairs of IgH and IgL genes were co-transfected in Expi293F cell culture. (B) Following expression, mAb were purified, and the purified IgG was subjected to SDS-PAGE analysis using a NuPAGE 4-12% Bis-Tris gel. Gel analysis revealed the presence of two bands in each lane, with approximate molecular weights of 25 kDa and 50 kDa, respectively. These bands corresponded to the light and heavy chains of the mAb and subjected to immunoblotting using the fusion proteins as indicated. To further confirm the identity of the heavy and light chains, Western blotting results displayed the heavy and light chains on the left side of the panel (lane P is positive control Human IgG 1K, lane 1 sample prepared under reducing conditions, lane 2 sample prepared under non-reducing conditions), confirming the successful expression and purification of the mAb. (C) For ELISA, purified Abs were diluted by serial dilution and incubated with purified IFN-α2a and IFN-β proteins. The reactivity of the purified Abs was determined by indirect ELISA, where the Ab titer was defined as the highest dilution of serum exhibiting an OD450 ratio. The results showed that the purified Abs exhibited high reactivity. The data are presented as the mean ± STDEV, with OD representing optical density. (D) In Western blot analysis, recombinant proteins (0.5 μg) were separated by SDS-PAGE, and equal concentrations of purified mAbs were used to probe the blots for purification and identification. A protein molecular weight marker (Lane M) was included. The primary Ab (at a dilution of 1:200) was recombinantly expressed with high specificity and affinity for the target proteins. The secondary Ab used was IRDye© 800 CW goat anti-human IgG (at a dilution of 1:10000). Western blotting confirmed that the observed band corresponded to the target proteins. Clones specific for IFN-α2a and IFN-α are indicated in orange fonts, while clones specific for IFN-β are indicated in purple fonts.

### Recombinant IgG production: Generation and characterization of IgG Abs

Sequenced DNA sequences described above encoding IgG heavy and light chains were synthetically constructed and inserted into the protein expression vector, pCDNA3.4, featuring a robust promoter (**Figure 2A**). Notably, the immediate early CMV promoter was employed to enhance mRNA stability and translation efficiency within the expression vector ^68^. Ab production was conducted in Expi293F cell culture, resulting in the successful expression of eight IgGs (6 target clones, BS-13H5, and CTI-AF1). Furthermore, LALA mutations were introduced to the AH07808, AH07824, AH07856, AH07857, AH07859, and AH07866 clones for additional validation. Purification of the recombinant proteins was achieved using a Ni-NTA agarose resin column, following the manufacturer’s instructions. Following expression, mAb were purified, and the purified IgG was subjected to SDS-PAGE analysis using a NuPAGE 4-12% Bis-Tris gel. The recombinant proteins expressed displayed a molecular weight of approximately 150 kDa under non-reducing conditions (**Figure 2B**, lane 2). Under denaturing conditions, the gel analysis revealed the presence of two bands in each lane with approximate molecular weights of 25 kDa and 50 kDa, corresponding to the light and heavy chains of the mAb. Confirmation of the heavy and light chains was also obtained by Western blotting to establish the successful expression and purification of each mAb (**Figure 2B**, lane 1).

The reactivity of expressed mAbs against recombinant IFN-α2a and IFN-β proteins was then evaluated using ELISA **(Figure 2C)** and Western blot analysis **(Figure 2D)**. Binding results demonstrated that clones AH07856 and AH07857 specifically reacted with IFN-α2a over AH07808, and all exhibited no cross-reactivity with IFN-β. Conversely, clones AH07859, AH07866, and AH07882 showed reactivity towards IFN-β over IFN-α by ELISA assays with minor detection of reactivity with IFN-α by Western for only AH07866 and AH07882. Added clones were generated (AH07824, AH07831, AH07845, AH07858, AH07881) yet showed binding to both IFN-α and IFN-β by either assay. Collectively, our findings confirm that human clones AH07856, AH07857, and AH07808 specifically recognize IFN-α2a without binding to IFN-β, while clones AH07859, AH07866, and AH07882 selectively recognize IFN-β with minimal to no cross-reactivity towards IFN-α2a.

### Identification of clones with neutralization anti-IFN-specific activity against either IFN-α- or IFN-β-mediated dimerization of the IFN receptor

We determined the neutralization potential of the top 10 identified clones (i.e. AH07808, AH07824, AH07831, AH07845, AH07856, AH07857, AH07858, AH07859, AH07866, AH07882) when evaluated for inhibition of IFN-α or IFN-β-mediated dimerization of the IFN receptor.

Positive control conditions using described neutralization Abs against IFN-α (BMS-13H5) and IFN-β (CTI-AF1) reconfirmed sensitivity of assay to detect IFN-specific inhibition (**Figure 3A**). Importantly, clones AH07856 and AH07857 inhibited IFN-α but not IFN-β, while clones AH07859 and AH07866 inhibited IFN-β but not IFN-α. These results were also confirmed for clones when the LALA mutation was introduced (**data not shown**).

**Figure 3.**
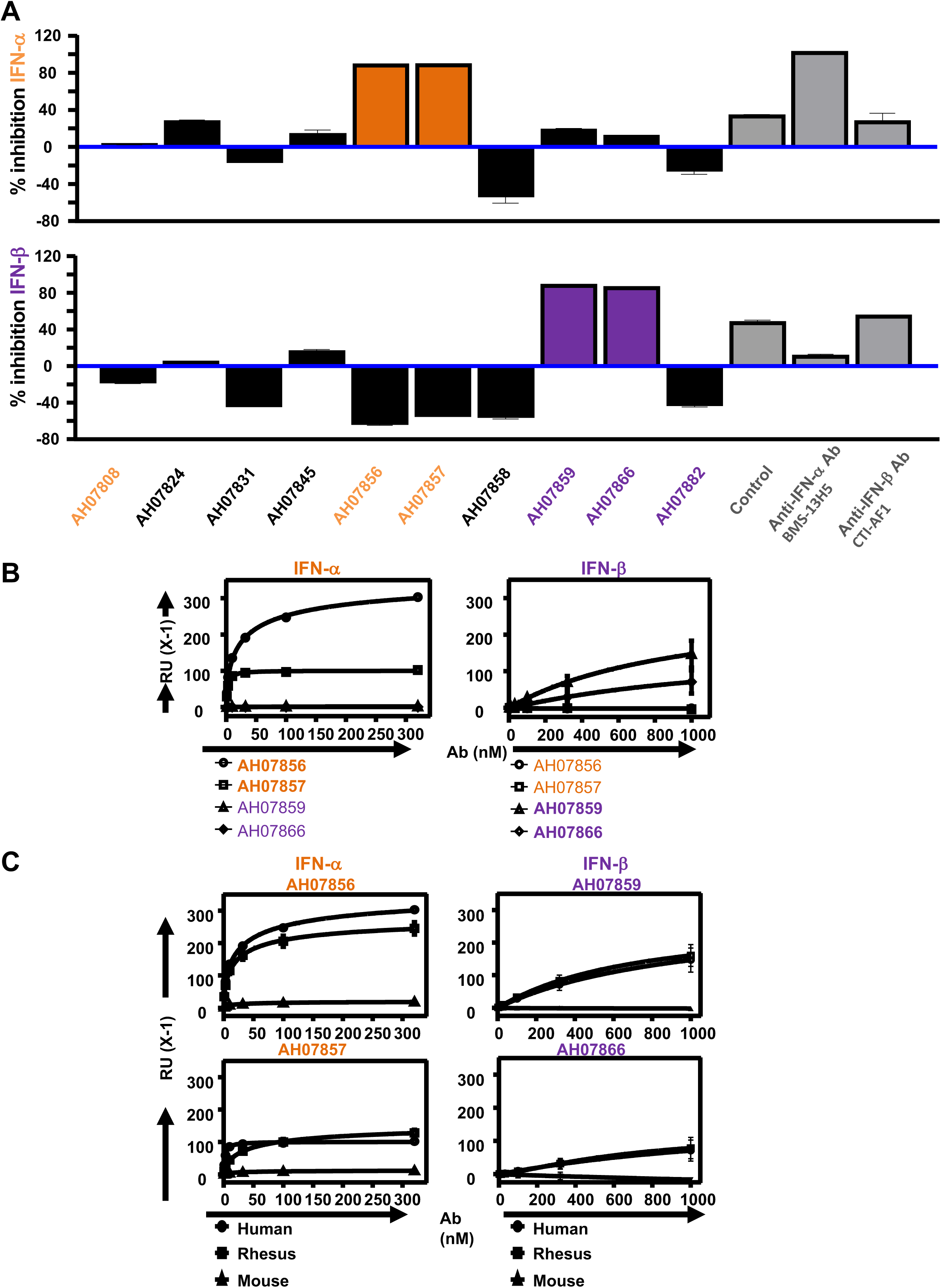
IFN-specific neutralization and binding saturation to human, rhesus or mouse IFN-α and IFN-β for anti-IFN-specific clones. (A) Percent (%) inhibition of IFNAR dimerization in the presence of IFN-α (top panel) or IFN-β (bottom panel) when combined with supernatants from IFN-specific binding clones, control plasma (MS-006 or Peg-008 from Fig 1, respectively), or neutralizing control anti-IFN-α (BMS-13H5) and anti-IFN-β (CTI-AF1) Abs. Supernatants from clones or anti-IFN-α or anti-IFN-β Abs were pre-incubated overnight with IFN-α or IFN-β at 4°C at the final concentrations. Final concentrations in the wells with HEK IFNAR1/IFNAR2 cells: 67% clone’s supernatants, 2.4 nM IFN-α, 1.2 nM IFN-β, 10 μg/ml of anti-IFN-α (BMS-13H5) and anti-IFN-β (CTI-AF1) Abs. (B) Saturation binding of different concentrations of neutralizing anti-IFN-specific clones AH07856, AH07857, AH07859 and AH07866 to human IFN-α (left) or human IFN-β (right) panels. (C) Left column shows saturation binding of different concentrations of anti-IFN-α-specific clones AH07856 (top) and AH07857 (bottom) to human, rhesus, and mouse IFN-α. Right column shows saturation binding of different concentrations of anti-IFN-β-specific clones AH07859 (top) and AH07866 (bottom) to human, rhesus, and mouse IFN-β. See **Figure S3** for added kinetic parameters for IFN-α and IFN-β clones binding to human, mouse and rhesus IFN-α and IFN-β, respectively. In panel (A) IFN-α and IFN-α-specific clones are indicated with orange fonts and orange shaded boxes respectively, while IFN-β and IFN-β-specific clones are indicated with purple fonts and purples shaded boxes respectively. In panels (B) and (C) saturation binding of different concentrations of clones to IFN-α and IFN-β from the different species are indicated with different shapes, while IFN-α and IFN-α-specific clones are indicated with orange fonts, and IFN-β and IFN-β-specific clones are indicated with purple fonts.

### Assessment of the binding affinity of the cloned anti-IFN-α and anti-IFN-β to human, rhesus and mouse IFN-α or IFN-β

To further explore the dimerization assay results, clones AH07856, AH07857, AH07859, and AH07866 were tested for their ability to bind either human IFN-α or IFN-β by surface plasmon resonance (SPR). The binding results for clones AH07856 and AH07857 are summarized in **Figure 3B** and **Table 1**. Briefly, clones AH07856 and AH07857 bound to IFN-α with nanomolar potency. However, no binding was detected by AH07859 or AH07866. By contrast, clones AH07859 and AH07866 were able to bind IFN-β, but no binding was detected for AH07856 or AH07857.

**Table 1.**
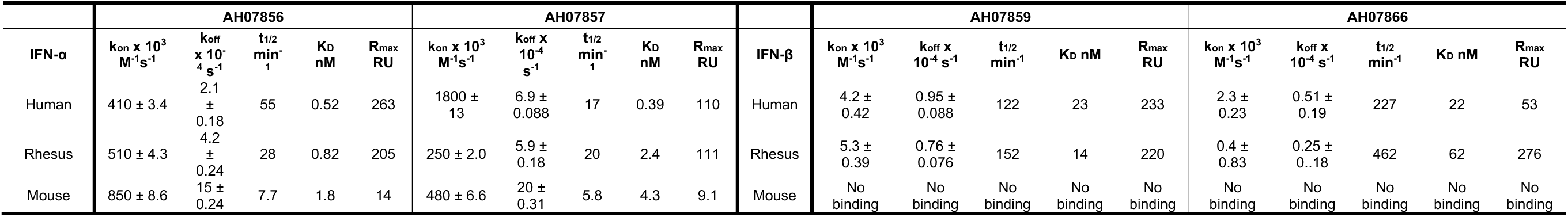
Binding kinetics of clones AH07856, AH07857, AH07859 and AH07866 with human, rhesus and mouse IFN-α2a and IFN-β analyzed by SPR.

Detailed kinetic studies were then carried out on these clones for their ability to bind human, rhesus, or mouse IFN-α or IFN-β (**Figure 3C, Figure S3, Table 1**). Briefly, the IFN-α clone AH07856 was able to bind both human and rhesus IFN-α with similar affinity (Kd values of 0.52 nM and 0.82 nM respectively), and also mouse IFN-α with slightly reduced affinity at 1.8 nM. The dissociation rate for AH07856 from mouse IFN-α was considerably faster compared to human and rhesus. Alternatively, the IFN-α clone AH07857 had much higher affinity for the human IFN-α (KD = 0.39 nM) compared to rhesus IFN-α (KD = 2.4 nM), or mouse IFN-α (KD = 4.3 nM). In addition, the dissociation rate for AH07857 from the human IFN-α was considerably slower compared to the rhesus and mouse IFN-α. By contrast, the IFN-β clones AH07859 and AH07866 were only able to bind both human and rhesus IFN-β, with no observable binding to mouse IFN-β. In addition, the affinities for these mAbs to IFN-β were considerably less than what was observed with the IFN-α mAbs, with KD values of 23 nM and 14 nM to human and rhesus IFN-β, respectively for AH07859, and 22 nM and 62 nM to human and rhesus IFN-β for AH07866. This weak affinity was caused by their apparent slow association rate together with even slower dissociation rates than what was observed for the IFN-α mAbs.

### Inhibition of IFN-mediated STAT1 phosphorylation in human cells by neutralizing anti-IFN-specific clones

To determine the ability of anti-IFN-specific clones AH07856, AH07857, AH07859 and AH07866 Abs to neutralize IFN-α or IFN-β signaling in primary lymphocytes, we first tested the induction of STAT-1 phosphorylation in primary CD4^+^ T cells by 1250 U/ml IFN-α or IFN-β in the presence of limiting dilutions of each cloned Ab starting at 25 μg/ml. Second, we tested the activity of each Ab at 25 μg/ml against decreasing amounts of IFN-α or IFN-β stimulation.

As expected by prior data, clones AH07856 and AH07857 strongly inhibited IFN-α but not IFN-β-mediated STAT-1 phosphorylation in CD4^+^ T cells (**Figure 4A, Figure S5**). Inhibition of pSTAT-1 phosphorylation was further confirmed for both anti-IFN-α-specific clones at all decreasing IFN-α concentrations tested. (**Figure S4**).

**Figure 4.**
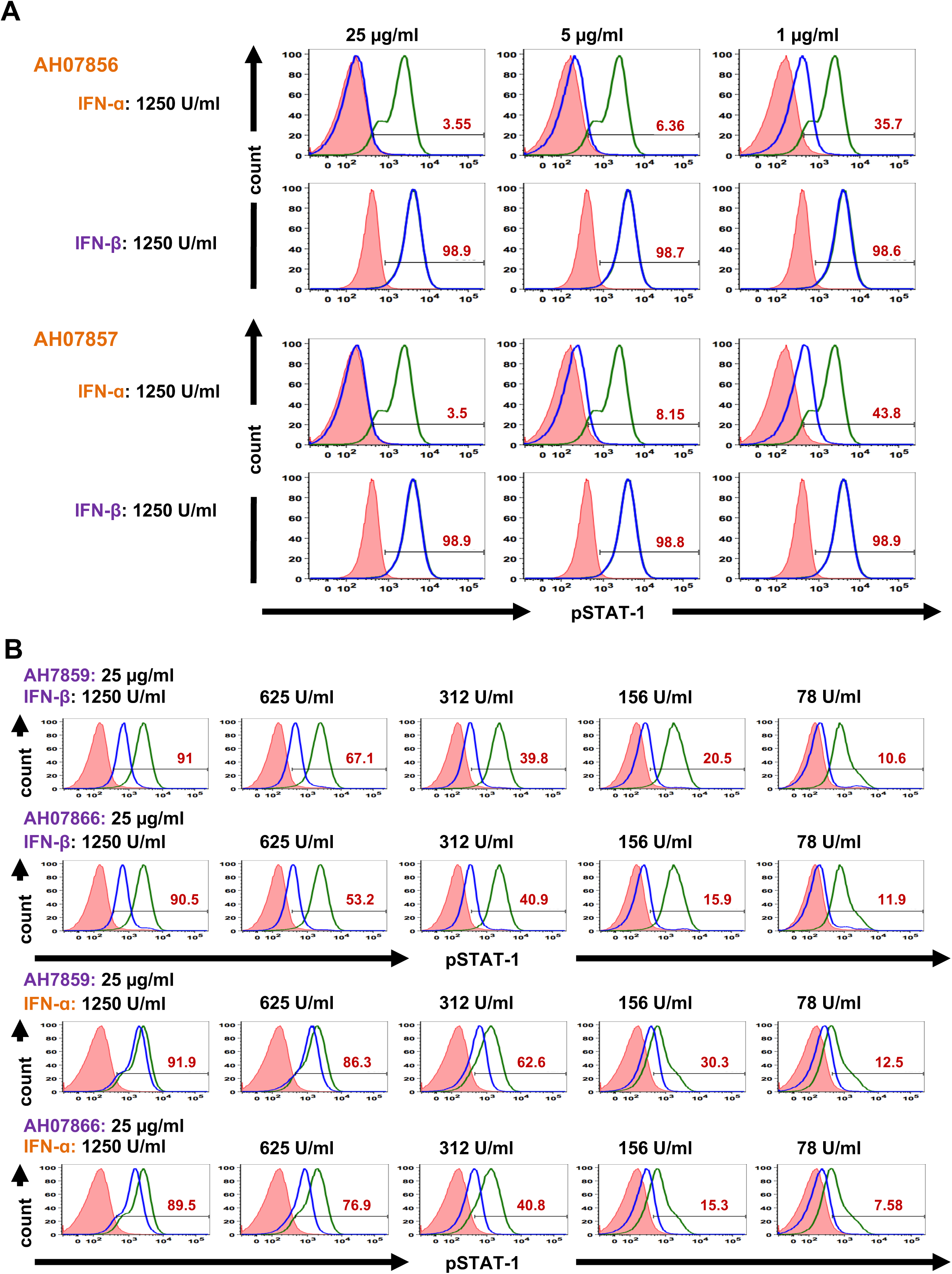
Inhibition of IFN-mediated STAT-1 phosphorylation in CD3^+^CD4^+^ primary T cells by anti-IFN-specific clones. Levels of phosphorylated STAT-1 (pSTAT-1) are shown in CD3^+^CD4^+^ T cells following stimulation of PBMC with or without IFN-α or IFN-β and in the presence or absence of anti-IFN-specific clones. (A) pSTAT-1 histogram panels for the effects of clones AH07856, or AH07857 (tested at 25, 5 and 1 ug/ml) on 1250 U/ml IFN-α or IFN-β stimulation. (B) pSTAT-1 histogram panels for the effects of clones AH7859 and AH07866 (tested at 25 μg/ml) on decreasing concentrations of IFN-α or IFN-β stimulation (1250, 625, 312, 156, 78 U/ml). For all histograms: (i) pink shaded peak shows constitutive pSTAT-1 levels in the absence of stimulation with IFN-α or IFN-β and in the absence of clones, (ii) green peak shows pSTAT-1 levels in the presence of stimulation with IFN-α or IFN-β and in the absence of clones, (iii) blue peak shows pSTAT-1 in CD3^+^CD4^+^ T cells in the presence of stimulation with IFN-α or IFN-β and in the presence of clones, and (iv) black line and number with red font inside the histogram show CD3^+^CD4^+^pSTAT-1^+^ percent (%) of CD3^+^CD4^+^ T cells following stimulation with IFN-α or IFN-β and in the presence of clones. IFN-α and IFN-α-specific clones are indicated with orange fonts, while IFN-β and IFN-β-specific clones are indicated with purple fonts. See **Figure S7** for gating strategy for the detection of pSTAT-1 in CD3^+^CD4^+^ T cells by flow cytometry.

On the other hand, clones AH07859 and AH07866 when tested at 25 μg/ml showed greater inhibition at lower concentrations of IFN-β (156 and 78U/ml) in contrast to partial inhibition observed at highest concentration of IFN-β tested (1250 U/ml) indicating greater potency with lower IFN-β levels (**Figure 4B**). Interestingly, usage in the assay of decreasing concentration of clones AH07859 and AH07866 following stimulation with 312 U/ml or lower IFN-α showed partial inhibition of IFN-α-mediated STAT-1 phosphorylation whereas IFN-β-mediated STAT-1 phosphorylation was more efficiently inhibited (**Figure 4B**). Overall, we confirm anti-IFN-specific inhibition of pSTAT-1 phosphorylation in primary cells exposed to either IFN-α-specific and anti-IFN-β-specific clones.

### Specificity of neutralizing anti-IFN-specific clones in inhibiting IFN-mediated gene expression in human myeloid cells following exposure to IFN-α subtypes or IFN-β

As STAT-1 phosphorylation is an acute inhibition effect, we further tested the ability of these Ab clones to inhibit pSTAT1 phosphorylation (30 mins after stimulation) and expression of IFN-induced genes MX1 and USP18 (24 hrs later) in PMA-differentiated THP-1 cells. We first titrated the inhibitory activity of the four anti-IFN-specific Abs when measuring MX1 induction by Western analysis at 24 hrs after IFN stimulation (**Figure 5A**). As before, we document IFN-specific inhibition of MX1 induction by all clones with greater potency by anti-IFN-α clones following IFN-α2a stimulation when compared to anti-IFN-β clones stimulated with IFN-β1a. Second, we tested by titration and breath the potency of the anti-IFN-α-specific clones when tested against 13 available IFN-α subtypes (**Table S1**) in addition to IFN-β1a. Both anti-IFN-α-specific clones effectively inhibited STAT-1 phosphorylation, MX1 and USP-18 by IFN-α2a, IFN-α2b, and IFN-αK (IFN-α6) with no inhibition activity against other IFN-α subtypes tested or IFN-β1a (**Figure 5B, Figure S6)**. Regarding anti-IFN-β-specific clones, we tested titrations of each clone (AH07859 and AH07866) or their combination against IFN-β1a-induced pSTAT-1, MX1 and UPS18 expression as above (**Figure 5C, D**). Similar inhibition was observed by either Ab used singly or by their combination confirming decreased IFN-β-stimulated ISG expression. Taken together, data supports that each identified neutralizing IFN-specific Ab against exogenous IFN-α or IFN-β can inhibit activation of ISG induction in T cells and in myeloid cells.

**Figure 5.**
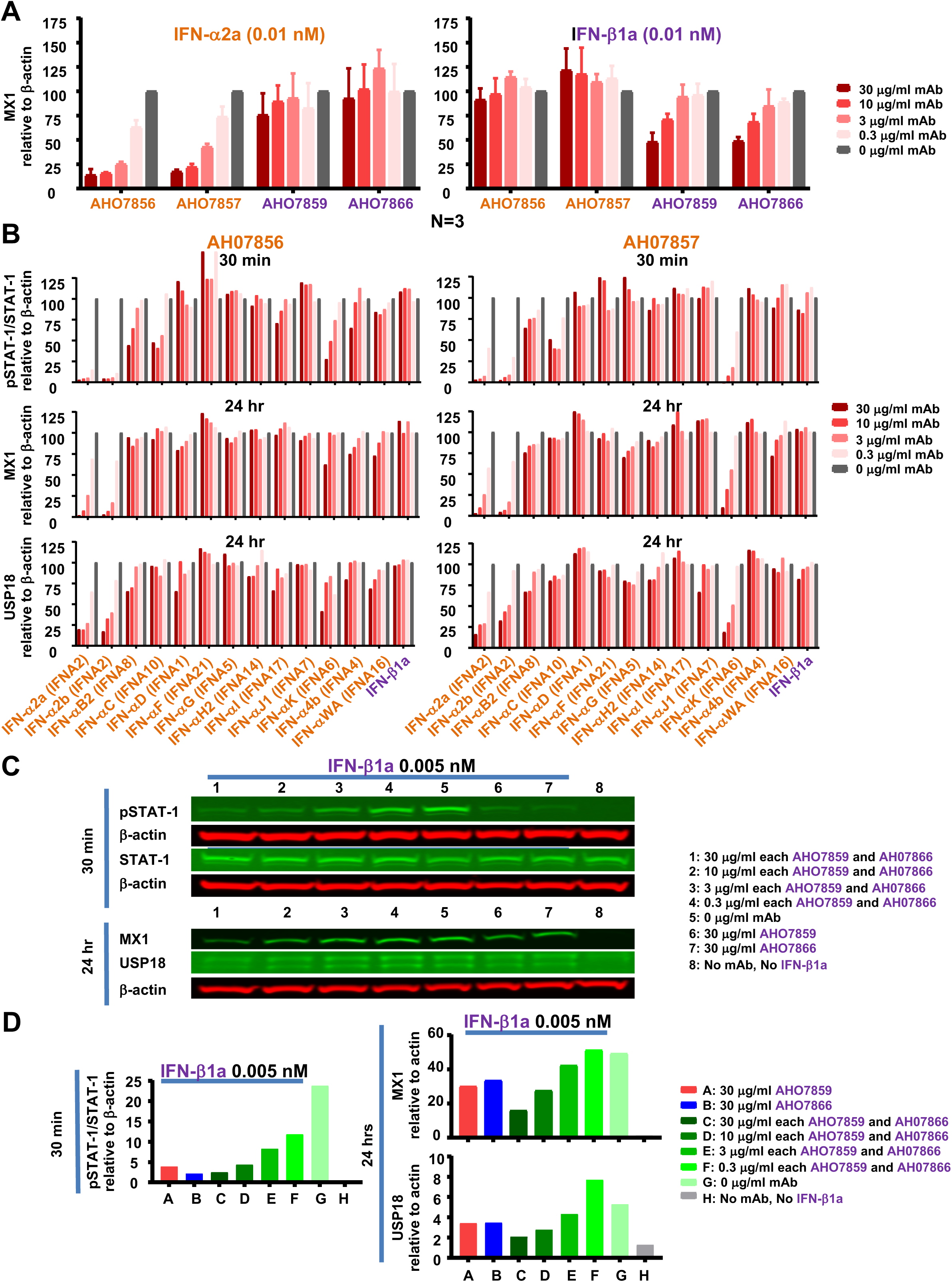
Inhibition of IFN-mediated expression of pSTAT-1, MX1 and USP18 in THP-1 cells by neutralizing anti-IFN-specific Abs. Shown are Western blots analysis for the expression of pSTAT-1, STAT-1, MX1, USP18 and β-actin in THP-1 cells following stimulation for 30 mins for pSTAT-1 and STAT-1, and for 24 hrs for MX1 and USP18 after IFN-α subtypes or IFN-β stimulation. (A) Bar graphs showing mean and standard error of 3 Western blots experiments measuring MX1 expression relative to β-actin levels in PMA-differentiated THP-1 cells following stimulation for 24 hrs with 0.01 nM of IFN-α2a or IFN-β1a in the presence or absence of clones AH07856, AH07857, AH07859 and AH07866 at 30, 10, 3, and 0.3 μg/ml. (B) Bar graphs showing mean and standard error of 3 Western blots experiments measuring pSTAT-1/STAT-1, MX1 and USP18 expression relative to β-actin levels in PMA-differentiated THP-1 cells following stimulation for 30 mins for STAT-1, and 24 hrs for MX1 and USP18 with 0.01 nM of IFN-α subtypes or IFN-β1a in the presence or absence of clones AH07856, and AH07857 at 30, 10, 3, and 0.3 μg/ml. Representative Western blots for these experiments are shown in **Figure S6**. (C) Western blots showing expression of pSTAT-1, STAT-1, MX1, USP18 and β-actin in PMA-differentiated THP-1 cells following stimulation for 30 mins for pSTAT-1 and STAT-1, and for 24 hrs for MX1 and USP18 after exposure to 0.005 nM of IFN-β1a in the presence or absence of clones AH07859 and AH07866 used alone or in combination at 30, 10, 3, and 0.3 μg/ml. (D) Bar graphs showing pSTAT-1/STAT-1, MX1 and USP18 expression relative to β-actin levels in PMA-differentiated THP-1 for the Western blots shown in panel (C). Western products evaluated were pSTAT-1/STAT-1: 91, 84kDa, MX1: 76 kDa, USP18: 39, 34 kDa, and β-actin: 42 kDa. In all panels, IFN-α, IFN-α subtypes and IFN-α-specific clones are indicated with orange fonts, while IFN-β, IFN-β1a and IFN-β-specific clones are indicated with purple fonts.

### Inhibition of type I IFN paracrine effects following LPS or S100A14 stimulation by neutralizing anti-IFN-specific Abs

To assess the neutralizing activity of the cloned anti-IFN-α-specific or anti-IFN-β-specific Abs against cells producing type IFNs following TLR stimulation, we tested the paracrine effects of TLR-4-induced type I IFN expression by LPS or S100A14 stimulation of PMA-differentiated THP1 cells. As shown in the top panel of **Figure 6**, STAT-1 phosphorylation was achieved after 30 mins stimulation of the cells with exogenously added IFN-α or IFN-β (**Figure 6 sections C and D**) with inhibition by clone AH07856 (**Figure 6 section C lane 1**) and AH07866 (**Figure 6 section D lane 2**) respectively. As expected, no detectable STAT-1 phosphorylation was observed at 30 mins after stimulation with LPS (**Figure 6 section A**) or S100A14 (**Figure 6 section B**) due to absence of signaling at this time otherwise present by induction of a prevalent IFN-β signature by 24 hrs ^67^ as evidenced by the induction ISG proteins MX1 and USP18. Evidence for the inhibition of paracrine effects of IFN-β in mediating ISG expression are indicated by the decreased levels of MX1 and USP18 observed at 24 hrs in the presence of neutralizing anti-IFN-β-specific (**Figure 6 sections A and B lane 2**) over anti-IFN-α-specific Ab (**Figure 6 sections A and B lane 1**). Importantly, the strength of neutralizing anti-IFN-α-specific and the neutralization activity of anti-IFN-β-specific Abs support their biological regulation potential.

**Figure 6.**
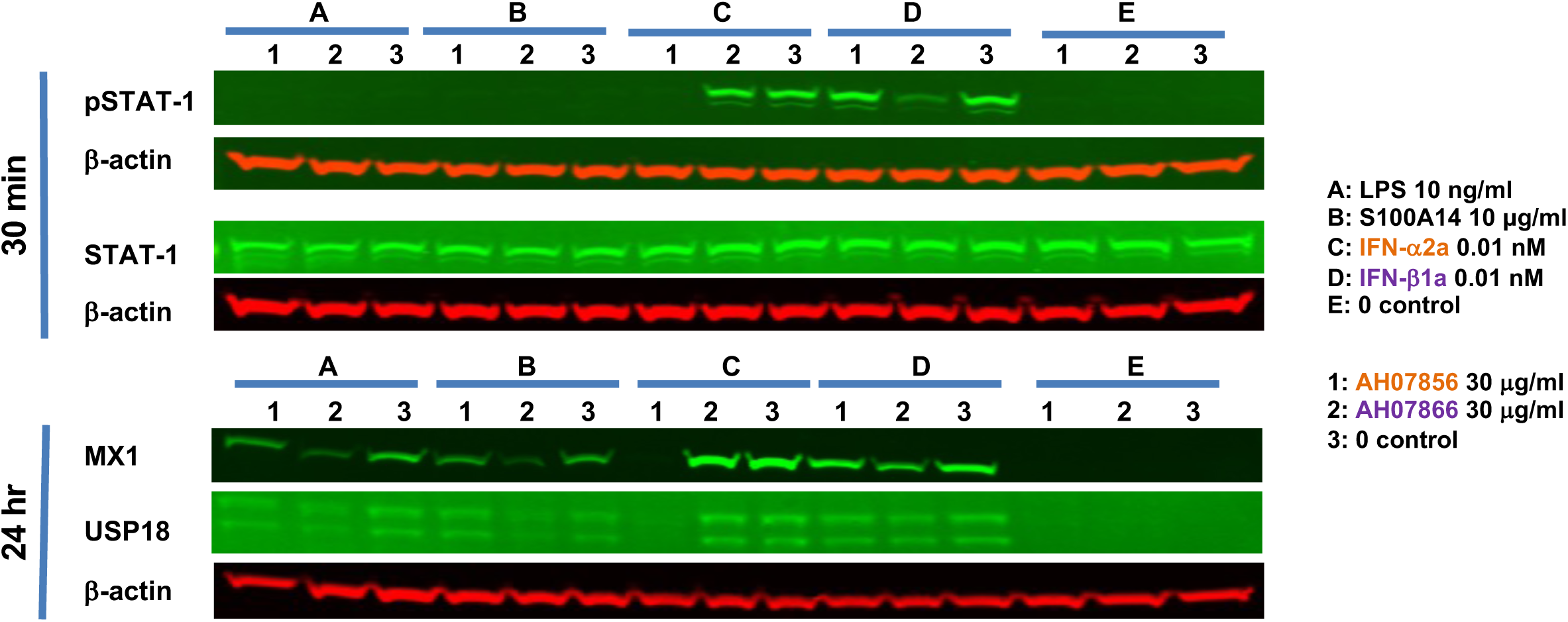
Inhibition of LPS or S100A14-induced IFN-mediated expression of MX1 and USP18 in THP-1 cells by neutralizing anti-IFN-specific Ab. Western blots showing the expression of pSTAT-1, STAT-1, MX1, USP18 and β-actin in THP-1 cells following stimulation for 30 mins for pSTAT-1 and STAT-1, and for 24 hrs for MX1 and USP18 with IFN-α2a (0.01 nM, section C), IFN-β1a (0.01 nM, section D), LPS (10 ng/ml, section A) or S100A14 (10 ng/ml, section B) or unstimulated control (section E) in the presence of 30 μg/ml of clones AH07856 (lane 1) or AH07866 (lane 2) or in the absence of clones (lane 3). pSTAT-1/STAT-1: 91, 84 kDa, MX1: 76 kDa, USP18: 39, 34 kDa, and β-actin: 42 kDa. IFN-α and IFN-α-specific clones are indicated with orange fonts, while IFN-β and IFN-β-specific clones are indicated with purple fonts.

## DISCUSSION

In this study we report novel natural human cloned neutralizing anti-IFN-specific Abs against IFN-α or IFN-β, able to modulate IFN-mediated signaling and gene expression. Importantly, we report success employing a phage-display strategy for cloning Abs using a pooled approach of cDNA from isolated total PBMC from multiple donors receiving IFN-α or IFN-β immunotherapy against HIV or to treat MS respectively, with pre-identified plasma Abs able to selectively neutralize IFN-α or IFN-β. By contrast to what was done in this study, conventional approaches to clone human mAbs are fraught with technical difficulties as they require the immortalization of human B cells or relay in sorting isolated antigen-specific B cells, and thus are not suitable for developing large panels of potentially therapeutic mAbs from target patients at once. Our data support that this strategy can be beneficial under the following conditions: (a) identification of patients with confirmed target plasma IgG-mediated activity, (b) availability of recombinant antigens for selective panning for phage selection, and (c) established functional assays able to evaluate and select target clones. The clinical impact of developing human neutralizing IFN-specific Abs against IFN-α and IFN-β is already evidenced by the clinical testing of the available humanized murine Abs targeting these molecules, as well as by the larger set of studies blocking all type I IFN signaling by directly targeting the IFNR ^66,69,70^. We now add two newly characterized natural human neutralizing anti-IFN-α-specific (AH07856, AH07857) and two anti-IFN-β-specific AH07859 and AH07866) Abs allowing for both basic and clinical studies targeting one or both human IFNs. Interestingly, we document that both neutralizing anti-IFN-α-specific Abs can target IFN-α2a and IFN-α2b as the major subtypes produced by myeloid and dendritic cells, as well as the IFN-αK subtype produced by keratinocytes and implicated in cutaneous lupus erythematosus and skin photosensitivity ^71,72^.

A clear greater potency of the natural neutralizing anti-IFN-α-specific Abs over the anti-IFN-β-specific Abs cloned was detected in various *in vitro* response assays using recombinant IFNs (rIFN) proteins. We interpret these differences to be related to greater *in vitro* efficiency of IFN-β to bind the IFNR over IFN-α in presence of Abs, or a weaker slower on-rate interaction by cloned anti-IFN-β Abs when compared to those against IFN-α as suggested by kinetics of saturation binding data. Importantly, these differences did not prevent these clones from acting to neutralize endogenously produced IFN-β levels following TLR activation as evidenced by lower ISG induction 24hrs later. Indeed, we establish the activity of anti-IFN-β-specific clone AH07866 to result in a greater inhibition of natural IFN-mediated effects in THP-1 cells following stimulation by either LPS, a major component of Gram-negative bacterial cell wall and a potent immunostimulatory product, or S100A14, a danger-associated molecular pattern and mediator of immune response ^73^. Both LPS and S100A14 are known to signal through Toll like receptor 4 (TLR4) ^74^ resulting in the production of type I IFNs ^75^. It is of interest to reconcile the “lower potency” interpretation of the anti-IFN-β Ab when tested with rIFN-β protein levels versus TLR- or S100A14 cell-stimulated levels of natural IFN-β suggesting rIFN levels used in biochemical or *in vitro* response assays may still represent an over-estimate of the physiologic target levels secreted by a cell. Our data also showed limited cross-reactivity by anti-IFN-β-specific Abs against IFN-α at lower concentrations as evaluated at STAT-1 induction 30 mins after stimulation, but without similar inhibition of ELISA binding, IFNR dimerization inhibition, MX1 induction 24 hours later, or detectable strength of binding affinity by SPR analysis, supporting the interpretation of a predominance of activity against IFN-β specificity over IFN-α.

Although our *in vitro* experiments document activity by all four Abs described, future studies will be needed to test their activity in vivo in order to reconfirm their potential to influence systemic IFN-mediated responses as predicted by our *in vitro* data. Regarding future in vivo testing, SPR analysis showed that all four neutralizing anti-IFN-specific clones had the ability to bind rhesus IFNs raising the possibility of testing a rhesus Ab version in this this model.

Type I IFNs have been shown to play a significant role in acute and chronic immune response and their regulation is evident in several clinical settings. As a result, the development of tools such as the Abs described here would allow to advance our understanding of type I IFN responses in cancer, autoimmune and infectious diseases ^1,76,77^ as exemplified by HIV infection where type I IFNs although antiviral have also been linked with lowering T cell mediated control of viral replication ^49–51,56^.

In conclusion, we present a novel approach for cloning neutralizing Abs against IFN-α and IFN-β from pooled peripheral blood mononuclear cells (PBMCs). Utilizing molecular and immunological techniques, variable gene transcripts for both heavy (VH) and light (VL) chains of IgGs were amplified. Two pairs of potent natural mAbs capable of selectively neutralizing either IFN-α or IFN-β were identified. These Abs hold promise for therapeutic strategies targeting autoimmune diseases, viral infections, and other medical applications. Further research is essential to fully elucidate the therapeutic potential of these Abs in preclinical and clinical settings.

## ACKNOWLEDGMENTS

We thank the persons who participated in the study.

## AUTHOR CONTRIBUTIONS

I.O., M.F., L.L, J.C. and S.M. performed experimental work. K.M, J.R.K., P.T., A.B-O recruited the patients and provided the samples. E.P., K.M., and L.J.M designed the study, evaluated the results, and wrote the manuscript. All participants have read and approved the manuscript.

## STAR★Methods

### KEY RESOURCES TABLE

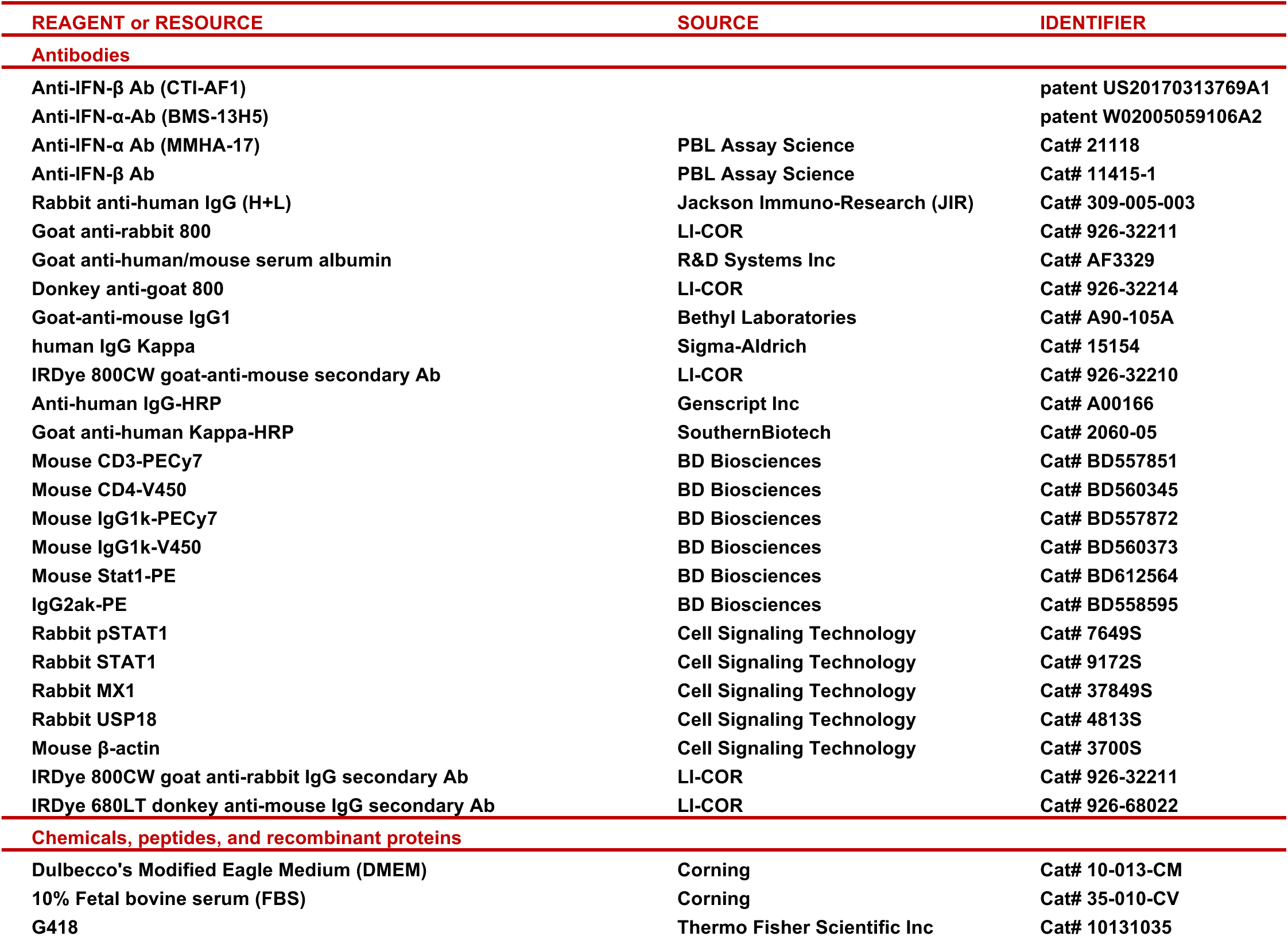
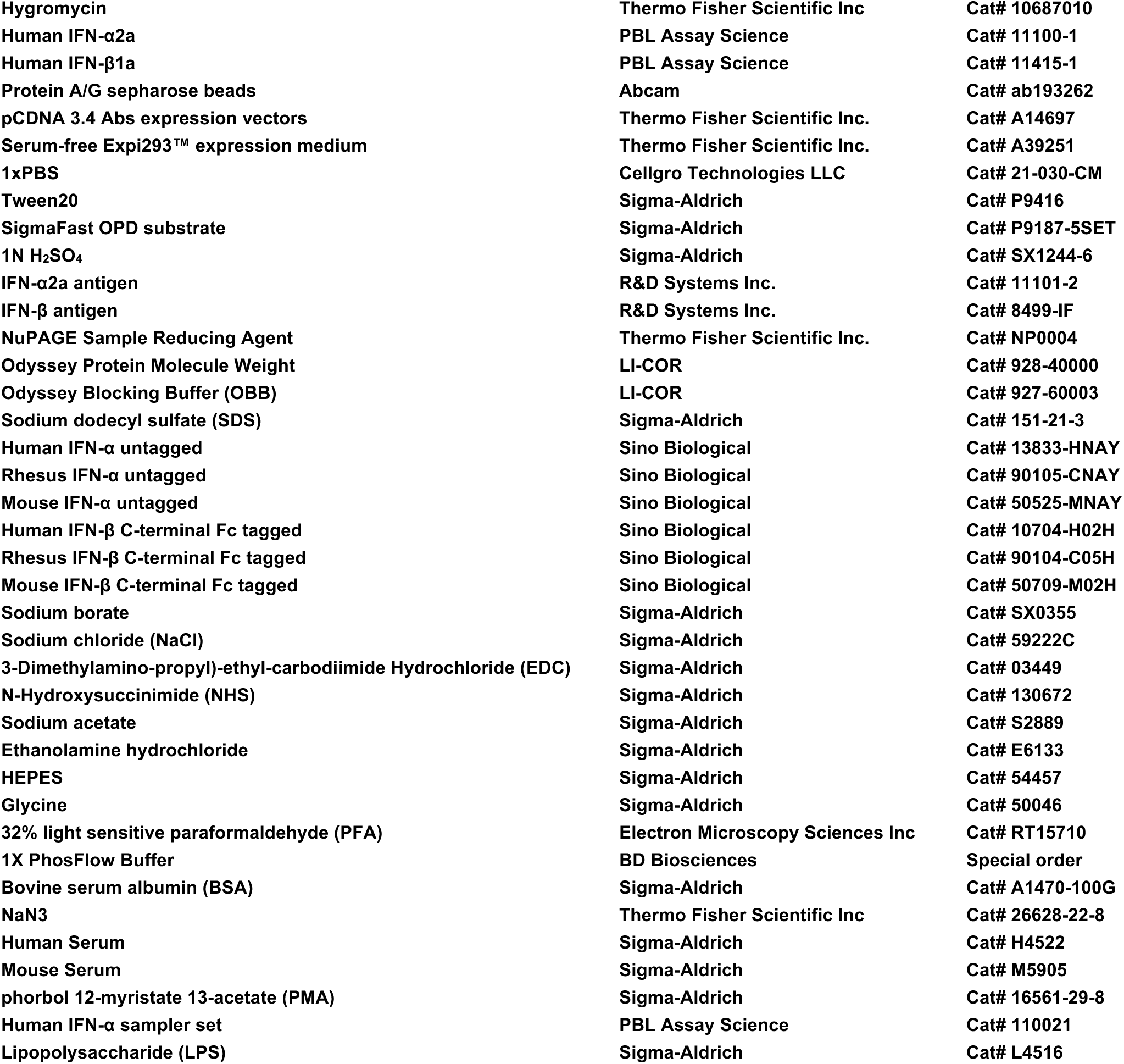
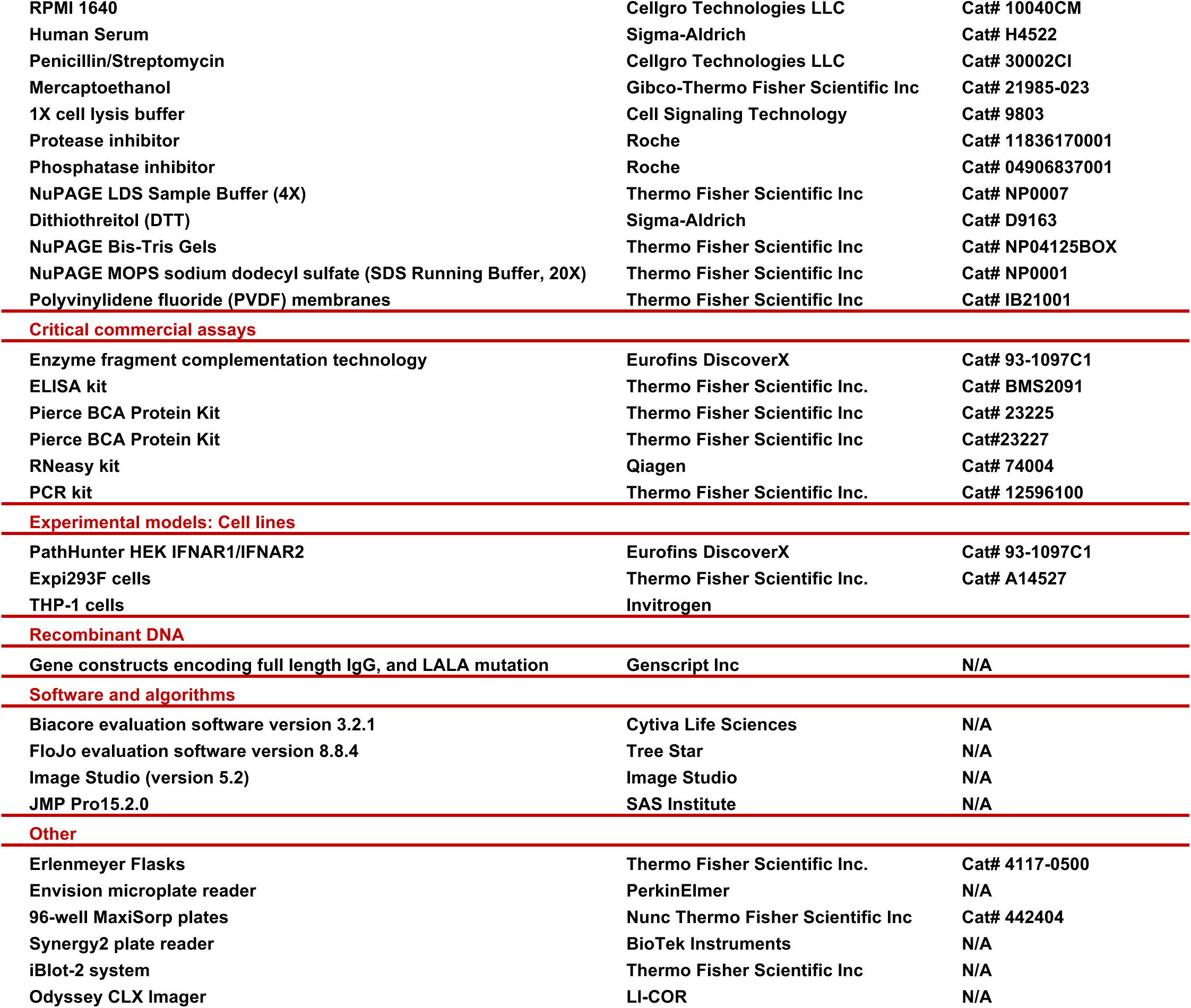
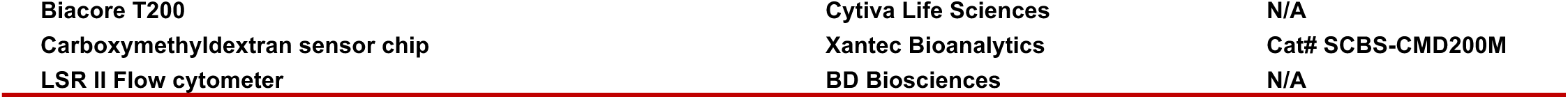

### RESOURCE AVAILABILITY

#### Lead contact

Further information and requests for resources and reagents should be directed to and will be fulfilled by the lead contact, Luis J. Montaner.

#### Materials availability

There are restrictions to the availability of antibodies (Abs) described due to the need to execute a material transfer agreement and availability of materials due to the lack of an external centralized repository for its distribution and our need to produce Abs. Upon MTA agreement, we are glad to share Abs with reasonable compensation by requestor for Ab generation, processing and shipping.

#### Data and code availability

- Data reported in this work paper are available from the lead contact upon request and executed data usage agreement.
- This paper does not report original code.
- Any additional information required to reanalyze the data reported in this work paper is available from the lead contact upon request.

## EXPERIMENTAL MODEL AND STUDY PARTICIPANT DETAILS

### Human subjects

Cryopreserved plasma from 11 participants with MS (**Table S2**), and from 18 persons living with HIV (**Table S2**) was screened for presence of anti-IFN-α and anti-IFN-β Abs respectively. The persons living with HIV were on suppressive ART (<50 HIV-1 copies/ml for ≥1 year, and ≥450 CD4^+^ T cells/mm^3^ at study entry) and received IFN-α as immunotherapy for 7.25 months under a research protocol NCT00594880 study ^28^, while the patients with MS were treated with IFN-β for a maximum of 6 months. A total of 220×10^6^ and 346×10^6^ cryopreserved PBMCs from two selected patients (MS-001 and Peg-008 respectively) with activity against IFN-β or IFN-α respectively were used for the cloning of anti-IFN-β and anti-IFN-α Abs as described below. The study protocol and informed consent procedures were approved by the Institutional Review Boards of the authors’ institutions.

### Cell lines

HEK IFNAR1/IFNAR2 cell line was from Eurofins DiscoverX. Expi293F cells were from Thermo Fisher Scientific Inc. THP-1 cells were from Invivogen (San Diego, CA). HEK IFNAR1/IFNAR2 cells were incubated at 37°C in Dulbecco’s Modified Eagle Medium (DMEM; Corning, New York, NY) containing 10% Fetal bovine serum (FBS; Corning), 0.8 mg/ml G418 (Thermo Fisher Scientific Inc, Philadelphia, PA), and 0.2 mg/ml Hygromycin (Thermo Fisher Scientific Inc). Expi293F cells were grown in serum-free Expi293™ expression medium (Thermo Fisher Scientific Inc) and maintained in Erlenmeyer Flasks (Thermo Fisher Scientific Inc) at 37°C with 8% CO_2_ on an orbital shaker. THP1 cells were cultured at 37°C with 8% CO_2_ in RPMI 1640 (Cellgro Technologies LLC, Lincoln, NE), containing 5% FBS (Corning), 1% Penicillin (Cellgro Technologies LLC), and 1% Streptomycin (Cellgro Technologies LLC).

## METHOD DETAILS

### Receptor dimerization assay

The presence of Abs against IFN-α or IFN-β was assessed using the IFN receptor dimerization assay. In this assay, IFN receptor dimerization is detected upon IFN binding in an engineered human embryonic kidney (HEK) cell line (HEK IFNAR1/IFNAR2) using enzyme fragment complementation technology (Eurofins DiscoverX, Fremont, CA). This cell line co-expresses IFNAR1 and IFNAR2, each conjugated to a fragment of β-galactosidase. Induction of receptor dimerization by IFN-α or IFN-β brings the enzyme fragments together to form an active enzyme, which hydrolyzes its substrate and generates a detectable chemiluminescence signal (Eurofins DiscoverX). The presence of anti-IFN-α or anti-IFN-β Abs, which bind to IFN-α or IFN-β, respectively, prevents receptor dimerization and the subsequent generation of chemiluminescence signal. HEK IFNAR1/IFNAR2 cells were maintained in Dulbecco’s Modified Eagle Medium (DMEM; Corning, New York, NY) containing 10% Fetal bovine serum (FBS; Corning), 0.8 mg/ml G418 (Thermo Fisher Scientific Inc, Philadelphia, PA), and 0.2 mg/ml Hygromycin (Thermo Fisher Scientific Inc). Briefly, cells were seeded in half-area white 96-well microplates (20000 cells/well) and incubated at 37°C overnight. Meanwhile, plasma samples or cloned anti-IFN-α or anti-IFN-β Abs were pre-incubated with human IFN-α2a (IFN-α, PBL Assay Science, Piscataway, NJ) or human IFN-β1a (IFN-β, PBL Assay Science) at 4°C overnight. For IFN-β, plasma was pre-incubated overnight with IFN-β at 4°C at 3X of the final concentrations, while for IFN-α, plasma was first heat inactivated at 56°C for 1 hr and then pre-incubated overnight with IFN-α at 4°C, at 3X of the final concentrations. Final concentrations in the wells with HEK IFNAR1/IFNAR2 cells were as shown: 11% plasma, or 67% clone supernatant, 2.4 nM IFN-α, and 1.2 nM IFN-β. After the overnight incubation, plasma pre-incubated with IFN-α or IFN-β was added to the cells and allowed to incubate at 37°C for 24 hrs. Added controls included plasma samples from two healthy individuals, F623-C32 and M486-B55 as negative controls, as well as expressed/purified humanized IgG CTI-AF1 (anti-IFN-β Ab, 10 μg/ml; patent US20170313769A1) or BMS-13H5 (neutralizing anti-IFN-α-specific Ab, 1.1 μg/ml or 10 μg/ml; patent W02005059106A2) ^66,67^ as positive controls, respectively. Luminescence signal was measured using the Envision microplate reader (PerkinElmer, Waltham, MA). The identification of samples with anti-IFN-α or anti-IFN-β Abs was based on the inhibition of IFN receptor dimerization despite the presence of IFN-α or IFN-β. Samples were run repeatedly in duplicates.

### Ab depletion

For Ab depletion, plasma sample at 10% in complete media was incubated with Protein A/G sepharose beads (Abcam, Boston, MA) for 1 hr with rotation at 4°C, for 5 consecutive times. For the receptor dimerization assay, the Ab depleted plasma samples were pre-incubated with IFN-β or IFN-α at 4°C overnight. Subsequently, HEK IFNAR1/IFNAR2 cell media was replaced by the Ab depleted plasma pre-incubated with IFN-β or IFN-α and incubated at 37°C for 24 hrs (final concentrations: 10% plasma, 1 nM IFN-β or 2.4 nM IFN-α). Luminescence signal was measured in a microplate reader. Detection of depletion of anti-IFN-β and anti-IFN-α Abs in the plasma sample by Western blotting was also performed. To detect Ab in the plasma sample, rabbit anti-human IgG (H+L) [Jackson Immuno-Research (JIR), Baltimore, PA] at 1:5000 was used as the primary Ab, and goat anti-rabbit 800 (LI-COR, Lincoln, NE) at 1:5000 as the secondary Ab. Albumin detection in the serum samples after consecutive rounds of Ab depletion was used to ensure the depletion was Ab specific. Following membrane stripping goat anti-human/mouse serum albumin (R&D Systems Inc, Minneapolis, MN) at 1:2500 was used as the primary Ab, and donkey anti-goat 800 (LI-COR) at 1:5000 as the secondary Ab.

### IFN-mAb development and recombinant mAb generation

Total RNA was isolated from PBMC samples using RNeasy kit (Qiagen, Germantown, MD) and pooled. After reverse transcription from pooled RNA using a reverse transcription polymerase chain reaction (PCR) kit (Thermo Fisher Scientific Inc), a master cDNA preparation was synthesized as a template to develop human Fab phagemid library. The VH and Fab-VL (Vk or Vl) genes were amplified by PCR followed by nested PCR using mixtures of primers specific for IgG and Igk and Igl. Amplified products from first-round and nested primer sets were then sequenced or IgG-VH and IgG-VL.

### Recombinant Plasmid Preparation and Ab expression

Recombinant Abs were cloned, as previously described ^78^. To generate recombinant Abs, H chain and L chain fragments were cloned into pCDNA 3.4 Abs expression vectors (Thermo Fisher Scientific Inc) in frame with either human IgG (Igk) constant domain.

For expression in mammalian cell lines, Ab constructs were first cloned into cytomegalovirus (CMV) expression vectors which relies on the widely utilized human CMV intermediate-early promoter/enhancer to drive overexpression of cloned inserts. Gene constructs encoding full length IgG, and LALA mutation are designed *in silico* and obtained from Genscript Inc (Piscataway, NJ) offering gene synthesis services. Transfection grade plasmids were maxi-prepared, for Expi293F cell expression (Thermo Fisher Scientific Inc). Expi293F cells (100 ml) were grown in serum-free Expi293™ expression medium (Thermo Fisher Scientific Inc). Following large-scale preparation of plasmid DNA, we used a suspension adapted HEK293 cell line (Expi293F) for high levels of transient expression. Optimization of heavy-to-light chain transfection ratios (IgG and Fab formats) maximized recombinant Ab yields. Cells were maintained in Erlenmeyer Flasks at 37°C with 8% CO_2_ on an orbital shaker. One day before transfection, the cells were seeded at an appropriate density. On the day of transfection, DNA and transfection reagent were mixed at an optimal ratio and then added into the flask with cells ready for transfection. Recombinant plasmids encoding target Ab were transiently co-transfected into Expi293F cell cultures. Cell culture supernatants collected on day 6 were used for Ab purification.

### Enzyme-linked immunosorbent assay (ELISA) and Western blot for hybridomas screening

ELISA was carried out using the 96-well MaxiSorp plates (Nunc Thermo Fisher Scientific Inc) coated with 1 µg/ml (volume 100 μl/well) of human recombinant IFN specific proteins (R&D Systems) in Phosphate Buffered Saline (PBS), pH 7.4 and incubated at 4°C overnight. Primary Abs were added at 1:50 dilution. Following incubation, plates were washed with PBS-T [1xPBS (Cellgro Technologies LLC, Lincoln, NE) with 0.05% Tween 20 (Sigma-Aldrich, St Louis, MO)] and blocked using PBS (Cellgro Technologies LLC) containing 10% FBS (Corning) for 1 hr at room temperature (RT). Subsequently, the plates were washed with PBS-T and incubated with hybridoma, serially diluted in PBS (Cellgro Technologies LLC) with 1% FBS (Corning) and 0.1% Tween 20 (Sigma-Aldrich) for 30 min on a shaker and 90 minutes at RT. After another wash, the plates were treated with goat-anti-mouse IgG1 (Bethyl Laboratories, Montgomery, TX) at a dilution of 1:10000 at RT for 1 hr. Post final wash, the plates were developed with SigmaFast OPD substrate (Sigma-Aldrich) for 5-10 min in the dark, and the reaction was stopped using 1 N H_2_SO_4_. (Sigma-Aldrich). The plates were read using a Synergy2 plate reader (BioTek Instruments, Winooski, VT) at an optical density (OD) of 450nm ^78^.

For the avidity test, the in-house avidity assay was standardized using a commercial ELISA kit (Thermo Fisher Scientific Inc) for detecting specific IgG Abs modified to incorporate an elution step with urea to remove low-avidity Abs from target antigen. For the assay, 100 µl of each serum diluted were added to wells of polystyrene plates coated with human IFN-α2a and IFN-β antigens (R&D Systems Inc). All serum samples were run twice in duplicate, as described before ^78^.

Western analysis was used to evaluate detection of IFNs and to analyze the light/heavy chain composition of cloned Ab preparations. For IFN detection, human recombinant IFN-α2a (1 µg/ml) and IFN-β (1 µg/ml) proteins (R&D Systems) were reduced using NuPAGE Sample Reducing Agent (10x) (Thermo Fisher Scientific Inc) and heating at 70°C for 10 min, then loaded onto sample lanes with Odyssey Protein Molecule Weight (LI-COR) serving as a standard marker. For cloned Ab composition evaluation, the mAbs following expression were purified and subjected to SDS-PAGE analysis using a NuPAGE 4-12% Bis-Tris gel (Thermo Scientific Inc) under reducing (lane 1) and non-reducing (lane 2) conditions. For Western analysis, an added positive human IgG Kappa (Sigma-Aldrich) loaded control lane was included (lane P). The gel electrophoresis was carried out using sodium dodecyl sulfate-12% polyacrylamide gel for 50 min at 200 V. Following electrophoresis, samples were transferred onto polyvinylidene fluoride (PVDF) membranes via an iBlot-2 system (Thermo Fisher Scientific Inc) and blocked using Odyssey Blocking Buffer (OBB) (LI-COR) for 1-2 hrs on a rocker. For IFN detection gels, membranes were treated with cloned Ab supernatant (1:200) in OBB (LI-COR) containing 0.1% Tween 20 (Sigma-Aldrich) at 4°C overnight. Following incubation, the membranes were washed four times at 5 min intervals with PBS-T. Subsequently, washed membranes were treated with IRDye 800 CW goat-anti-human secondary Ab (LI-COR) in OBB (LI-COR) containing 0.1% Tween 20 (Sigma-Aldrich) and 0.01% SDS (Sigma-Aldrich) at a dilution of 1:10000 and incubated at RT for 60 min in the dark on the rocker. For Ab clone composition evaluations, primary Ab goat anti-human IgG-HRP (GenScript Inc), and goat anti-human Kappa-HRP (SouthernBiotech, Birmingham, AL) were used. Following incubation, the membranes were rewashed four times and scanned using Odyssey CLX Imager (LI-COR).

### Assessment of the binding of the cloned anti-IFN-α or anti-IFN-β Abs on IFN-α or IFN-β by surface plasmon resonance

IFN Abs affinity was assessed by surface plasmon resonance (SPR) using a Biacore T200 (Cytiva Life Sciences, Marlborough, MA). Approximately, 300 response units (RU) of IFN-α (human, rhesus, or mouse untagged, Sino Biological, Chesterbrook, PA) or 500 RU of IFN-β (human, rhesus, or mouse, C-terminal Fc tagged, Sino Biological) was immobilized on carboxymethyldextran sensor chip (CMD200M, Xantec Bioanalytics, Germany) using amine coupling. Briefly, the sensor chip was first washed with 0.1 M sodium borate, pH 9.0 (Sigma-Aldrich), 1 M NaCl (Sigma-Aldrich) for 3 min at 10 ml/min, followed by activation with 50 mM 3-Dimethylamino-propyl)-ethyl-carbodiimide Hydrochloride (EDC)/100 mM N-Hydroxysuccinimide (NHS) (Sigma-Aldrich) for 12 min at 10 ml/min. Either IFN-α or IFN-β in 10 mM sodium acetate, pH 5.0 (Sigma-Aldrich) was then flowed over the chip until the desired immobilization level was achieved. After a 30 min delay, the remaining activated sites were blocked with 1 M ethanolamine hydrochloride, pH 8.5 (Sigma-Aldrich) for 5 min at 10 ml/min. The running buffer was then switched from distilled water to 10 mM HEPES, pH 7.4 (Sigma-Aldrich), 150 mM NaCl (Sigma-Aldrich), 0.1% Tween20 (Sigma-Aldrich). Antibodies were serially diluted 1:3.16 for 6 concentrations starting at 320 nM for IFN-α Abs, and 1000 nM for IFN-β Abs. The association and dissociation times were 180/420 sec for IFN-α Abs, and 360/600 sec for IFN-β Abs at 25 ml/min. After the respective dissociation time, the Ab was dissociated by a 60 sec injection of 20 mM glycine, pH 2.0 (Sigma-Aldrich) at 30 ml/min. Kinetic parameters were obtained from global nonlinear regression fits of the data using Biacore evaluation software (Cytiva Life Sciences).

### Assessment of the effect of the cloned anti-IFN-α or anti-IFN-β Abs on the IFN-α or IFN-β-mediated STAT-1 phosphorylation in primary cells by flow cytometry

Two million PBMC (10×10^6^ cells/ml) were added into 15 ml conical tubes and stained with surface Abs (CD3-PECy7, CD4-V450) or corresponding surface isotypes (IgG1k-PECy7, IgG1k-V450) at 4°C for 30 min. Cells were then washed with RT 1XPBS (Cellgro Technologies LLC) at 1500 rpm for 5 min and re-suspended in warm 1 ml 1XPBS Cellgro Technologies LLC). Subsequently, the cloned anti-IFN-α or anti-IFN-β Abs were incubated with IFN-α (PBL Assay Science) or IFN-β (PBL Assay Science) at RT for 15 min and then added to the cells at 37°C for 10 min. Cells were then fixed with 16% light sensitive paraformaldehyde (PFA, Electron Microscopy Sciences Inc, Hatfield, PA) at a final concentration of 5% PFA at 37°C for 10 min. Subsequently, 10ml of 1XPBS were added and cells were centrifuged at 2200 rpm for 10 min. Supernatant was removed, the cell pellet was loosen up, and incubated with 500 µl – 1 ml of 1X PhosFlow Buffer (BD Biosciences, San Diego, CA, USA) at RT for 30 min. Then, 3 ml FACS wash [1xPBS (Cellgro Technologies LLC) supplemented with 0.1% BSA (Sigma-Aldrich) and 0.02% NaN_3_ (Thermo Fisher Scientific Inc), supplemented with 5% human serum (Sigma-Aldrich) and 5% mouse serum (Sigma-Aldrich)] was added, cells were centrifuged at 2200 rpm for 10 min, the FACS wash was removed, intracellular Stat1-PE or corresponding intracellular isotype (IgG2ak-PE was added in corresponding tubes and incubated in the dark at RT for 1 hr. Then, 3 ml FACS wash were added, cells were centrifuged at 2200 rpm for 10 min, FACS wash was removed, cells were transferred into Facs tubes and run on LSR II Flow cytometer (BD Biosciences). A minimum of 100000 events were collected in the live cell gate (FSC/SSC). Data were analyzed using FloJo software (Version 8.8.4, Tree Star, Ashland, OR). The gating strategy for analysis of STAT-1 expression in CD3+CD4+ cells is shown in **Figure S7**. All Abs were from BD Biosciences.

### Assessment of the effect of the cloned anti-IFN-α or anti-IFN-β Abs on the IFN-α or IFN-β-mediated STAT-1 phosphorylation, IFN-induced GTP-binding protein (MX-1) expression, and ubiquitin specific peptidase 18 (USP18) expression in THP-1 cells by Western blot

In order to assess the effect of cloned anti-IFN-α or anti-IFN-β Abs on IFN-α or IFN-β-mediated protein expression, THP-1 cells (Invivogen) were seeded in 6-well plates (1.5×10^6^ cells/well) with or without phorbol 12-myristate 13-acetate (PMA, 100 ng/ml, 2 ml/well, Sigma-Aldrich) and incubated at 37°C for 48 hrs. At the end of the 48 hrs incubation, cloned anti-IFN-α or anti-IFN-β Abs were pre-incubated with human IFN-α2a (different subtypes of human IFN-α (human IFN-α sampler set, PBL Assay Science), human IFN-β1a (IFN-β, PBL Assay Science), lipopolysaccharide (LPS, Sigma-Aldrich), or the S100 calcium binding protein A14 (S100A14, ACROBiosystems, Newark, DE) at RT for 30 min at RT, and then were added to the cells for an additional incubation at 37°C for 30 mins (for assessment of STAT-1 phosphorylation) or 24 hrs (for assessment of MX1 and USP18 expression). At the end of the incubation the RPMI culture media [RPMI 1640 (Cellgro Technologies LLC), 10% human serum (Sigma-Aldrich), 100 U/ml Penicillin (Cellgro Technologies LLC), 100 μg/ml Streptomycin (Cellgro Technologies LLC), 55 μM 2-Mercaptoethanol (Gibco-Thermo Fisher Sicientific)] was replaced with fresh RPMI culture media and cells were further incubated at 37°C for 8 hrs. Cells were subsequently washed with ice cold 1XPBS (Cellgro Technologies LLC) twice and then lysed with 150 μl 1X cell lysis buffer (Cell Signaling Technology, Danvers, MA), on ice for 10 min followed by removal of the lysed cells, sonication for 10 sec and centrifugation at 13000 g in a benchtop centrifuge at 4°C for 15 min. The protein concentration was determined in the samples using Pierce BCA Protein Kit (Thermo Fisher Scientific Inc) and 20 μg of protein from each sample were used for Western blot.

Briefly, for immunoblotting, cells were lysed with 1x cell lysis buffer (Cell Signaling Technology) containing protease inhibitor and phosphatase inhibitor (Roche, Basel, Switzerland). The lysates were incubated on ice for 10 min followed by removal of the lysed cells and sonication for 10 sec, and then centrifugation at 13000 g in a benchtop centrifuge at 4°C for 15 min. The protein concentration was determined in the samples using the Pierce BCA Protein Kit (Thermo Fisher Scientific Inc). The solid protein was denatured at 95°C for 5 min in NuPAGE LDS Sample Buffer (4X) (Thermo Scientific Inc) containing 50 mM dithiothreitol (DTT, Sigma-Aldrich). Samples were loaded to 4-12% NuPAGE Bis-Tris Gels (Thermo Scientific Inc) and run at 150 V for 1 hr with NuPAGE MOPS sodium dodecyl sulfate (SDS Running Buffer, 20X, Thermo Scientific Inc). Following electrophoresis, the samples were transferred onto PVDF membranes via an iBlot-2 system (Thermo Fisher Scientific Inc) and blocked using LI-COR Blocking Buffer (OBB, LI-COR) for 1-2 hrs on a rocker. The primary Abs were treated in OBB containing 0.2% Tween-20 overnight at 4°C on the rocker. The following primary Abs, used at 1:1000 dilution, were purchased from Cell Signaling Technology: pSTAT1, STAT1, MX1, USP18 and β-actin. The secondary Abs, IRDye 800 CW, and IRDye 680 LT, were used at 1:5000 dilution, were purchased from LI-COR). The Western blot images were recorded using the Odyssey CLX imager (LI-COR) and the Western blot quantification was done using Image Studio (version 5.2).

## QUANTIFICATION AND STATISTICAL ANALYSIS

Data were summarized as means and standard deviation (STDEV) using JMP Pro15.2.0 (SAS Institute, Cary, NC). In the dimerization assays, means and STDEV of percent (%) inhibition was used, which was calculated by considering the luminescence value of the control sample without IFN (“no IFN”) as 100 % inhibition and the luminescence value of the control sample with IFN (“IFN only”) as 0% inhibition and by using the following formula: 100 – [(sample – “no IFN”) / (“IFN only” – “no IFN”) * 100]. In Western data means and STDEV of normalized data from 3 experiments were calculated. Western data for each experiment were analyzed using Imager Studio (version 5.2) and first normalized to β-actin equal control for each sample, and then normalized to either no-stimulated control, or IFN-α or IFN-β alone without Ab control. The no-stimulation control was 100%, while the stimulation over the no-stimulation control was above 100%.

## SUPPLEMENTAL FIGURE TITLES AND LEGENDS

**Figure S1. Specificity of Ab depletion in plasma from participant MS-001**

Left panel shows western blots of 10% plasma from participant MS-001 following 5 rounds of depletion of Abs with A/G sepharose beads. Middle and right panels show the same blot following stripping before incubation with anti-albumin Ab, and after incubation with anti-albumin Ab respectively.

**Figure S2. A flowchart summarizing the generation of antigen-specific mAbs against IFN-α2a and IFN-β from peripheral PBMCs, and unrooted phylogenetic tree of the identified binding Abs**

(A) Cognate pairs of VH and VL genes were amplified by PCR and used as shown in the flowchart.

(B-C) Unrooted phylogenetic tree was constructed based on CDR comparisons of sequences from anti-IFN-α2a and IFN-β Abs to evaluate relatedness. Included in analysis were sequences to previously reported neutralizing anti-IFN-α (BMS-13H5) or anti-IFN-β (CTI-AF1) Abs respectively.

**Figure S3. Kinetic parameters for IFN-α clones AH07856 and AH07857, as well as for IFN-β clones AH07859 and AH07866 binding to human, mouse and rhesus IFN-α and IFN-β respectively**

(A) Binding kinetics of clones AH07856 (top panel) and AH07857 (bottom panel) over time are shown for different concentrations of human, rhesus, and mouse IFN-α using different colors of lines.

(B) Binding kinetics of clones AH07859 (top panel) and AH07866 (bottom panel) over time are shown for different concentrations of human, rhesus, and mouse IFN-β using different colors of lines.

In panel (A) human, rhesus, and mouse IFN-α and IFN-α-specific clones are indicated with orange fonts, while in panel (B) human, rhesus, and mouse IFN-β and IFN-β-specific clones are indicated with purple fonts.

**Figure S4. Inhibition of IFN-mediated STAT-1 phosphorylation in CD3^+^CD4^+^ primary T cells by anti-IFN-α-specific clones when cells are stimulated with decreasing concentrations of IFN-α but not when cells are stimulated with decreasing concentrations of IFN-β**

Levels of phosphorylated STAT-1 (pSTAT-1) are shown in CD3^+^CD4^+^ T cells following stimulation of PBMC with or without IFN-α or IFN-β (in decreasing concentrations) and in the presence or absence of anti-IFN-α-specific clones.

(A) pSTAT-1 histogram panels for the effects of clones AH07856, or AH07857 (tested at 25 μg/ml) on decreasing concentrations of IFN-α stimulation (1250, 625, 312, 156, 78 U/ml).

(B) pSTAT-1 histogram panels for the effects of clones AH07856, or AH07857 (tested at 25 μg/ml) on decreasing concentrations of IFN-β stimulation (1250, 625, 312, 156, 78 U/ml). For all histograms: (i) pink shaded peak shows constitutive pSTAT-1 levels in the absence of stimulation with IFN-α or IFN-β and in the absence of clones, (ii) green peak shows pSTAT-1 levels in the presence of stimulation with IFN-α or IFN-β and in the absence of clones, (iii) blue peak shows pSTAT-1 in CD3^+^CD4^+^ T cells in the presence of stimulation with IFN-α or IFN-β and in the presence of clones, and (iv) black line and number with red font inside the histogram show CD3^+^CD4^+^pSTAT-1^+^ percent (%) of CD3^+^CD4^+^ T cells following stimulation with IFN-α or IFN-β and in the presence of clones. IFN-α and IFN-α-specific clones are indicated with orange fonts, while IFN-β and IFN-β-specific clones are indicated with purple fonts. See **Figure S7** for gating strategy for the detection of pSTAT-1 in CD3^+^CD4^+^ T cells by flow cytometry.

**Figure S5. Increased inhibition of IFN-β-mediated STAT-1 phosphorylation in CD3^+^CD4^+^ T cells by anti-IFN-β-specific clones following cell stimulation with lower concentration of IFN-β**

Levels of phosphorylated STAT-1 (pSTAT-1) are shown in CD3^+^CD4^+^ T cells following stimulation of PBMC with or without IFN-α or IFN-β and in the presence or absence of anti-IFN-specific clones.

(A) pSTAT-1 histogram panels for the effects of clones AH07859, or AH07866 (tested at 25, 5, 1 μg/ml) following stimulation with 1250 U/ml of IFN-α or IFN-β.

(B) pSTAT-1 histogram panels for the effects of clones AH07856, AH07857, AH07859, or AH07866 (tested at 25, 5, 1 μg/ml) following stimulation with 156 U/ml of IFN-α or IFN-β. For all histograms: (i) pink shaded peak shows constitutive pSTAT-1 levels in the absence of stimulation with IFN-α or IFN-β and in the absence of clones, (ii) green peak shows pSTAT-1 levels in the presence of stimulation with IFN-α or IFN-β and in the absence of clones, (iii) blue peak shows pSTAT-1 in CD3^+^CD4^+^ T cells in the presence of stimulation with IFN-α or IFN-β and in the presence of clones, and (iv) black line and number with red font inside the histogram show CD3^+^CD4^+^pSTAT-1^+^ percent (%) of CD3^+^CD4^+^ T cells following stimulation with IFN-α or IFN-β and in the presence of clones. IFN-α and IFN-α-specific clones are indicated with orange fonts, while IFN-β and IFN-β-specific clones are indicated with purple fonts. See **Figure S7** for gating strategy for the detection of pSTAT-1 in CD3^+^CD4^+^ T cells by flow cytometry.

**Figure S6. Inhibition of IFN-α2a-, IFN-α2b-, and IFN-αK-together with lack of inhibition of IFN-αI-, IFN-αJ1-, IFN-αD-, IFN-αF-, IFN-αB2- or IFN-αC-mediated STAT-1 phosphorylation, and MX1 and USP18 expression by clones AH07856 and AH07857 in PMA-differentiated THP-1 cells**

Western blots showing expression of pSTAT-1, STAT-1, MX1, USP18 and β-actin in PMA-differentiated THP-1 cells following stimulation for 30 mins for pSTAT-1 and STAT-1, and for 24 hrs for MX1 and USP18 with 0.01 nM of (A) IFN-α2a, IFN-αH2, IFN-α2b, or IFN-α2G, (B) IFN-αK, IFN-α4b, or IFN-αWA, (C) IFN-αI, IFN-αJ1, IFN-αD, or IFN-αF, and (D) IFN-αB2 or IFN-αC in the presence or absence of clones AH07856 and AH07857 at 30, 10, 3, and 0.3 μg/ml. pSTAT-1/STAT-1: 91, 84 kDa, MX1: 76 kDa, USP18: 39, 34 kDa, and β-actin: 42 kDa. IFN-α subtypes and IFN-α-specific clones are indicated with orange fonts, while IFN-β1a is indicated with purple fonts.

**Figure S7. Gating strategy for the detection of pSTAT-1 in CD3^+^CD4^+^ T cells by flow cytometry**

Shown is gating strategy starting with left panel gating by FSC-A versus SSC-A dot plot of all cells with gate isolating the lymphocytes. From lymphocyte gate, a CD4 versus CD3 dot plot is generated with gate isolating the CD3^+^CD4^+^ T cells. From CD3^+^CD4^+^ gate, a histogram for level of pSTAT-1 is generated with threshold gate added to delineate baseline expression in the absence of IFN stimulation. In summary, gating allows for identification of the frequency of CD3^+^CD4^+^pSTAT-1^+^ cells before and after IFN stimulation.

## SUPPLEMENTAL TABLE TITLES AND LEGENDS

**Table S1. Nomenclature and genes of human IFN-α**

**Table S2. Demographics of participants with MS and of persons living with HIV-1**

## REFERENCES

1. McNab, F., Mayer-Barber, K., Sher, A., Wack, A., and O’Garra, A. (2015). Type I interferons in infectious disease. Nat Rev Immunol 15, 87–103. 10.1038/nri3787.

2. Biron, C.A. (2001). Interferons alpha and beta as immune regulators--a new look. Immunity 14, 661–664. S1074-7613(01)00154-6 [pii].

3. Malmgaard, L. (2004). Induction and regulation of IFNs during viral infections. J Interferon Cytokine Res 24, 439–454. 10.1089/1079990041689665.

4. Stark, G.R., Kerr, I.M., Williams, B.R., Silverman, R.H., and Schreiber, R.D. (1998). How cells respond to interferons. Annu Rev Biochem 67, 227–264. 10.1146/annurev.biochem.67.1.227.

5. Asmuth, D.M., Utay, N.S., and Pollard, R.B. (2016). Peginterferon alpha-2a for the treatment of HIV infection. Expert Opin Investig Drugs 25, 249–257. 10.1517/13543784.2016.1132699.

6. de Weerd, N.A., Vivian, J.P., Nguyen, T.K., Mangan, N.E., Gould, J.A., Braniff, S.J., Zaker-Tabrizi, L., Fung, K.Y., Forster, S.C., Beddoe, T., et al. (2013). Structural basis of a unique interferon-beta signaling axis mediated via the receptor IFNAR1. Nat Immunol 14, 901–907. 10.1038/ni.2667.

7. Isaacs, A., and Lindenmann, J. (1957). Virus interference. I. The interferon. Proc R Soc Lond B Biol Sci 147, 258–267. 10.1098/rspb.1957.0048.

8. Bach, E.A., Aguet, M., and Schreiber, R.D. (1997). The IFN gamma receptor: a paradigm for cytokine receptor signaling. Annu Rev Immunol 15, 563–591. 10.1146/annurev.immunol.15.1.563.

9. Kotenko, S.V., Gallagher, G., Baurin, V.V., Lewis-Antes, A., Shen, M., Shah, N.K., Langer, J.A., Sheikh, F., Dickensheets, H., and Donnelly, R.P. (2003). IFN-lambdas mediate antiviral protection through a distinct class II cytokine receptor complex. Nat Immunol 4, 69–77. 10.1038/ni875.

10. Sheppard, P., Kindsvogel, W., Xu, W., Henderson, K., Schlutsmeyer, S., Whitmore, T.E., Kuestner, R., Garrigues, U., Birks, C., Roraback, J., et al. (2003). IL-28, IL-29 and their class II cytokine receptor IL-28R. Nat Immunol 4, 63–68. 10.1038/ni873.

11. Stanifer, M.L., Pervolaraki, K., and Boulant, S. (2019). Differential Regulation of Type I and Type III Interferon Signaling. Int J Mol Sci 20. 10.3390/ijms20061445.

12. Lee, A.J., and Ashkar, A.A. (2018). The Dual Nature of Type I and Type II Interferons. Front Immunol 9, 2061. 10.3389/fimmu.2018.02061.

13. Pestka, S., Krause, C.D., and Walter, M.R. (2004). Interferons, interferon-like cytokines, and their receptors. Immunol Rev 202, 8–32. 10.1111/j.0105-2896.2004.00204.x.

14. Gibbert, K., Schlaak, J.F., Yang, D., and Dittmer, U. (2013). IFN-alpha subtypes: distinct biological activities in anti-viral therapy. Br J Pharmacol 168, 1048–1058. 10.1111/bph.12010.

15. Schreiber, G. (2017). The molecular basis for differential type I interferon signaling. J Biol Chem 292, 7285–7294. 10.1074/jbc.R116.774562.

16. Kalie, E., Jaitin, D.A., Podoplelova, Y., Piehler, J., and Schreiber, G. (2008). The stability of the ternary interferon-receptor complex rather than the affinity to the individual subunits dictates differential biological activities. J Biol Chem 283, 32925–32936. 10.1074/jbc.M806019200.

17. Jaks, E., Gavutis, M., Uze, G., Martal, J., and Piehler, J. (2007). Differential receptor subunit affinities of type I interferons govern differential signal activation. J Mol Biol 366, 525–539. 10.1016/j.jmb.2006.11.053.

18. Platanias, L.C., and Fish, E.N. (1999). Signaling pathways activated by interferons. Exp Hematol 27, 1583–1592. 10.1016/s0301-472x(99)00109-5.

19. Goodbourn, S., Didcock, L., and Randall, R.E. (2000). Interferons: cell signalling, immune modulation, antiviral response and virus countermeasures. J Gen Virol 81, 2341–2364. 10.1099/0022-1317-81-10-2341.

20. Waddell, S.J., Popper, S.J., Rubins, K.H., Griffiths, M.J., Brown, P.O., Levin, M., and Relman, D.A. (2010). Dissecting interferon-induced transcriptional programs in human peripheral blood cells. PLoS One 5, e9753. 10.1371/journal.pone.0009753.

21. Dill, M.T., Makowska, Z., Trincucci, G., Gruber, A.J., Vogt, J.E., Filipowicz, M., Calabrese, D., Krol, I., Lau, D.T., Terracciano, L., et al. (2014). Pegylated IFN-alpha regulates hepatic gene expression through transient Jak/STAT activation. J Clin Invest 124, 1568–1581. 10.1172/JCI70408.

22. Marijanovic, Z., Ragimbeau, J., van der Heyden, J., Uze, G., and Pellegrini, S. (2007). Comparable potency of IFNalpha2 and IFNbeta on immediate JAK/STAT activation but differential down-regulation of IFNAR2. Biochem J 407, 141–151. 10.1042/BJ20070605.

23. Marcellin, P., Boyer, N., Gervais, A., Martinot, M., Pouteau, M., Castelnau, C., Kilani, A., Areias, J., Auperin, A., Benhamou, J.P., et al. (1997). Long-term histologic improvement and loss of detectable intrahepatic HCV RNA in patients with chronic hepatitis C and sustained response to interferon-alpha therapy. Ann Intern Med 127, 875–881. 10.7326/0003-4819-127-10-199711150-00003.

24. Lee, H.W., Lee, J.S., and Ahn, S.H. (2020). Hepatitis B Virus Cure: Targets and Future Therapies. Int J Mol Sci 22. 10.3390/ijms22010213.

25. Liang, T.J., Block, T.M., McMahon, B.J., Ghany, M.G., Urban, S., Guo, J.T., Locarnini, S., Zoulim, F., Chang, K.M., and Lok, A.S. (2015). Present and future therapies of hepatitis B: From discovery to cure. Hepatology 62, 1893–1908. 10.1002/hep.28025.

26. Doyle, T., Goujon, C., and Malim, M.H. (2015). HIV-1 and interferons: who’s interfering with whom? Nat Rev Microbiol 13, 403–413. 10.1038/nrmicro3449.

27. Lane, H.C., Davey, V., Kovacs, J.A., Feinberg, J., Metcalf, J.A., Herpin, B., Walker, R., Deyton, L., Davey, R.T., Jr., Falloon, J., and, et al. (1990). Interferon-alpha in patients with asymptomatic human immunodeficiency virus (HIV) infection. A randomized, placebo-controlled trial. Ann Intern Med 112, 805–811. 10.7326/0003-4819-112-11-805.

28. Azzoni, L., Foulkes, A.S., Papasavvas, E., Mexas, A.M., Lynn, K.M., Mounzer, K., Tebas, P., Jacobson, J.M., Frank, I., Busch, M.P., et al. (2013). Pegylated Interferon alfa-2a monotherapy results in suppression of HIV type 1 replication and decreased cell-associated HIV DNA integration. J Infect Dis 207, 213–222. 10.1093/infdis/jis663.

29. Papasavvas, E., Azzoni, L., Kossenkov, A.V., Dawany, N., Morales, K.H., Fair, M., Ross, B.N., Lynn, K., Mackiewicz, A., Mounzer, K., et al. (2019). NK Response Correlates with HIV Decrease in Pegylated IFN-alpha2a-Treated Antiretroviral Therapy-Suppressed Subjects. J Immunol 203, 705–717. 10.4049/jimmunol.1801511.

30. Papasavvas, E., Azzoni, L., Pagliuzza, A., Abdel-Mohsen, M., Ross, B.N., Fair, M., Howell, B., Hazuda, D., Chomont, N., Li, Q., et al. (2020). Safety, immune and anti-viral effects of pegylated interferon alpha 2b administration in ART-suppressed individuals: Results of pilot clinical trial. AIDS Res Hum Retroviruses. 10.1089/AID.2020.0243.

31. Sun, H., Buzon, M.J., Shaw, A., Berg, R.K., Yu, X.G., Ferrando-Martinez, S., Leal, M., Ruiz-Mateos, E., and Lichterfeld, M. (2014). Hepatitis C therapy with interferon-alpha and ribavirin reduces CD4 T-cell-associated HIV-1 DNA in HIV-1/hepatitis C virus-coinfected patients. J Infect Dis 209, 1315–1320 10.1093/infdis/jit628.

32. Angel, J.B., Greaves, W., Long, J., Ward, D., Rodriguez, A.E., Scevola, D., and DeJesus, E. (2009). Virologic and immunologic activity of PegIntron in HIV disease. AIDS 23, 2431–2438. 10.1097/QAD.0b013e32832f30ca.

33. de Jong, H.J.I., Kingwell, E., Shirani, A., Cohen Tervaert, J.W., Hupperts, R., Zhao, Y., Zhu, F., Evans, C., van der Kop, M.L., Traboulsee, A., et al. (2017). Evaluating the safety of beta-interferons in MS: A series of nested case-control studies. Neurology 88, 2310–2320. 10.1212/WNL.0000000000004037.

34. Pestka, S. (2007). The interferons: 50 years after their discovery, there is much more to learn. J Biol Chem 282, 20047–20051. 10.1074/jbc.R700004200.

35. Neutralizing antibodies during treatment of multiple sclerosis with interferon beta-1b: experience during the first three years. The IFNB Multiple Sclerosis Study Group and the University of British Columbia MS/MRI Analysis Group. (1996). Neurology 47, 889–894. 10.1212/wnl.47.4.889.

36. Sominanda, A., Lundkvist, M., Fogdell-Hahn, A., Hemmer, B., Hartung, H.P., Hillert, J., Menge, T., and Kieseier, B.C. (2010). Inhibition of endogenous interferon beta by neutralizing antibodies against recombinant interferon beta. Arch Neurol 67, 1095–1101. 10.1001/archneurol.2010.218.

37. Shapiro, A.M., Jack, C.S., Lapierre, Y., Arbour, N., Bar-Or, A., and Antel, J.P. (2006). Potential for interferon beta-induced serum antibodies in multiple sclerosis to inhibit endogenous interferon-regulated chemokine/cytokine responses within the central nervous system. Arch Neurol 63, 1296–1299. 10.1001/archneur.63.9.1296.

38. Hemmer, B., Stuve, O., Kieseier, B., Schellekens, H., and Hartung, H.P. (2005). Immune response to immunotherapy: the role of neutralising antibodies to interferon beta in the treatment of multiple sclerosis. Lancet Neurol 4, 403–412. 10.1016/S1474-4422(05)70117-4.

39. Sbardella, E., Tomassini, V., Gasperini, C., Bellomi, F., Cefaro, L.A., Morra, V.B., Antonelli, G., and Pozzilli, C. (2009). Neutralizing antibodies explain the poor clinical response to interferon beta in a small proportion of patients with multiple sclerosis: a retrospective study. BMC Neurol 9, 54. 10.1186/1471-2377-9-54.

40. Sorensen, P.S. (2008). Neutralizing antibodies against interferon-Beta. Ther Adv Neurol Disord 1, 125–141. 10.1177/1756285608095144.

41. Bekisz, J.B., zur Nedden, D.L., Enterline, J.C., and Zoon, K.C. (1989). Antibodies to interferon-alpha 2 in patients treated with interferon-alpha 2 for hairy cell leukemia. J Interferon Res 9 *Suppl 1*, S1–7.

42. Spiegel, R.J., Jacobs, S.L., and Treuhaft, M.W. (1989). Anti-interferon antibodies to interferon-alpha 2b: results of comparative assays and clinical perspective. J Interferon Res 9 *Suppl 1*, S17–24.

43. Russo, D., Candoni, A., Zuffa, E., Minisini, R., Silvestri, F., Fanin, R., Zaja, F., Martinelli, G., Tura, S., Botta, G., and Baccarani, M. (1996). Neutralizing anti-interferon-alpha antibodies and response to treatment in patients with Ph+ chronic myeloid leukaemia sequentially treated with recombinant (alpha 2a) and lymphoblastoid interferon-alpha. Br J Haematol 94, 300–305. 10.1046/j.1365-2141.1996.d01-1790.x.

44. Oberg, K., and Alm, G.V. (1989). Development of neutralizing interferon antibodies after treatment with recombinant interferon-alpha 2b in patients with malignant carcinoid tumors. J Interferon Res 9 *Suppl 1*, S45–49.

45. Itri, L.M., Sherman, M.I., Palleroni, A.V., Evans, L.M., Tran, L.L., Campion, M., and Chizzonite, R. (1989). Incidence and clinical significance of neutralizing antibodies in patients receiving recombinant interferon-alpha 2a. J Interferon Res 9 *Suppl 1*, S9–15.

46. Arai, K., Shindo, M., and Okuno, T. (1994). [Anti-interferon antibody in chronic hepatitis C]. Nihon Rinsho 52, 1929–1934.

47. Steinmann, G.G., God, B., Rosenkaimer, F., Adolf, G., Bidlingmaier, G., Fruhbeis, B., Lamche, H., Lindner, J., Patzelt, E., Schmahling, C., and, et al. (1992). Low incidence of antibody formation due to long-term interferon-alpha 2c treatment of cancer patients. Clin Investig 70, 136–141. 10.1007/BF00227355.

48. Scagnolari, C., Trombetti, S., Solda, A., Milella, M., Gaeta, G.B., Angarano, G., Scotto, G., Caporaso, N., Morisco, F., Cozzolongo, R., et al. (2012). Development and specificities of anti-interferon neutralizing antibodies in patients with chronic hepatitis C treated with pegylated interferon-alpha. Clin Microbiol Infect 18, 1033–1039. 10.1111/j.1469-0691.2011.03729.x.

49. Bialas, A.R., Presumey, J., Das, A., van der Poel, C.E., Lapchak, P.H., Mesin, L., Victora, G., Tsokos, G.C., Mawrin, C., Herbst, R., and Carroll, M.C. (2017). Microglia-dependent synapse loss in type I interferon-mediated lupus. Nature 546, 539–543. 10.1038/nature22821.

50. McGlasson, S., Jury, A., Jackson, A., and Hunt, D. (2015). Type I interferon dysregulation and neurological disease. Nat Rev Neurol 11, 515–523. 10.1038/nrneurol.2015.143.

51. Taylor, J.M., Moore, Z., Minter, M.R., and Crack, P.J. (2018). Type-I interferon pathway in neuroinflammation and neurodegeneration: focus on Alzheimer’s disease. J Neural Transm (Vienna) 125, 797–807. 10.1007/s00702-017-1745-4.

52. Dillon, S.M., Guo, K., Austin, G.L., Gianella, S., Engen, P.A., Mutlu, E.A., Losurdo, J., Swanson, G., Chakradeo, P., Keshavarzian, A., et al. (2018). A compartmentalized type I interferon response in the gut during chronic HIV-1 infection is associated with immunopathogenesis. AIDS 32, 1599–1611. 10.1097/QAD.0000000000001863.

53. Ng, C.T., Sullivan, B.M., Teijaro, J.R., Lee, A.M., Welch, M., Rice, S., Sheehan, K.C., Schreiber, R.D., and Oldstone, M.B. (2015). Blockade of interferon Beta, but not interferon alpha, signaling controls persistent viral infection. Cell Host Microbe 17, 653–661. 10.1016/j.chom.2015.04.005.

54. Dagenais-Lussier, X., Loucif, H., Murira, A., Laulhe, X., Stager, S., Lamarre, A., and van Grevenynghe, J. (2017). Sustained IFN-I Expression during Established Persistent Viral Infection: A "Bad Seed" for Protective Immunity. Viruses 10. 10.3390/v10010012.

55. Dianzani, F., Rozera, G., Abbate, I., D’Offizi, G., Abdeddaim, A., Vlassi, C., Antonucci, G., Narciso, P., Martini, F., and Capobianchi, M.R. (2008). Interferon may prevent HIV viral rebound after HAART interruption in HIV patients. J Interferon Cytokine Res 28, 1–3. 10.1089/jir.2007.0076.

56. Morimoto, Y., Kishida, T., Kotani, S.I., Takayama, K., and Mazda, O. (2018). Interferon-beta signal may up-regulate PD-L1 expression through IRF9-dependent and independent pathways in lung cancer cells. Biochem Biophys Res Commun 507, 330–336. 10.1016/j.bbrc.2018.11.035.

57. Carnathan, D., Lawson, B., Yu, J., Patel, K., Billingsley, J.M., Tharp, G.K., Delmas, O.M., Dawoud, R., Wilkinson, P., Nicolette, C., et al. (2018). Reduced Chronic Lymphocyte Activation following Interferon Alpha Blockade during the Acute Phase of Simian Immunodeficiency Virus Infection in Rhesus Macaques. J Virol 92. 10.1128/JVI.01760-17.

58. Swainson, L.A., Sharma, A.A., Ghneim, K., Ribeiro, S.P., Wilkinson, P., Dunham, R.M., Albright, R.G., Wong, S., Estes, J.D., Piatak, M., et al. (2022). IFN-alpha blockade during ART-treated SIV infection lowers tissue vDNA, rescues immune function, and improves overall health. JCI Insight 7. 10.1172/jci.insight.153046.

59. Teijaro, J.R., Ng, C., Lee, A.M., Sullivan, B.M., Sheehan, K.C., Welch, M., Schreiber, R.D., de la Torre, J.C., and Oldstone, M.B. (2013). Persistent LCMV infection is controlled by blockade of type I interferon signaling. Science 340, 207–211. 10.1126/science.1235214.

60. Zemek, R.M., Chin, W.L., Fear, V.S., Wylie, B., Casey, T.H., Forbes, C., Tilsed, C.M., Boon, L., Guo, B.B., Bosco, A., et al. (2022). Temporally restricted activation of IFNbeta signaling underlies response to immune checkpoint therapy in mice. Nat Commun 13, 4895. 10.1038/s41467-022-32567-8.

61. Yuliantie, E., Dai, X., Yang, D., Crack, P.J., and Wang, M.W. (2018). High-throughput screening for small molecule inhibitors of the type-I interferon signaling pathway. Acta Pharm Sin B 8, 889–899. 10.1016/j.apsb.2018.07.005.

62. Cornez, I., Yajnanarayana, S.P., Wolf, A.M., and Wolf, D. (2017). JAK/STAT disruption induces immuno-deficiency: Rationale for the development of JAK inhibitors as immunosuppressive drugs. Mol Cell Endocrinol 451, 88–96. 10.1016/j.mce.2017.01.035.

63. Kirou, K.A., and Gkrouzman, E. (2013). Anti-interferon alpha treatment in SLE. Clin Immunol 148, 303–312. 10.1016/j.clim.2013.02.013.

64. Chasset, F., and Arnaud, L. (2018). Targeting interferons and their pathways in systemic lupus erythematosus. Autoimmun Rev 17, 44–52. 10.1016/j.autrev.2017.11.009.

65. Schwartz, D.M., Kanno, Y., Villarino, A., Ward, M., Gadina, M., and O’Shea, J.J. (2017). JAK inhibition as a therapeutic strategy for immune and inflammatory diseases. Nat Rev Drug Discov 16, 843–862. 10.1038/nrd.2017.201.

66. Neelakantan, S., Oemar, B., Johnson, K., Rath, N., Salganik, M., Berman, G., Pelletier, K., Cox, L., Page, K., Messing, D., and Tarabar, S. (2021). Safety, Tolerability, and Pharmacokinetics of PF-06823859, an Anti-Interferon beta Monoclonal Antibody: A Randomized, Phase I, Single- and Multiple-Ascending-Dose Study. Clin Pharmacol Drug Dev 10, 307–316. 10.1002/cpdd.887.

67. Nistler, R., Sharma, A., Meeth, K., Huard, C., Loreth, C., Kalbasi, A., Tyminski, E., Bellmore, R., Coyle, A.J., Gulla, S.V., et al. (2020). TLR Stimulation Produces IFN-beta as the Primary Driver of IFN Signaling in Nonlymphoid Primary Human Cells. Immunohorizons 4, 332–338. 10.4049/immunohorizons.1800054.

68. Choi, H., Ho, M., Adeniji, O.S., Giron, L., Bordoloi, D., Kulkarni, A.J., Puchalt, A.P., Abdel-Mohsen, M., and Muthumani, K. (2021). Development of Siglec-9 Blocking Antibody to Enhance Anti-Tumor Immunity. Front Oncol 11, 778989. 10.3389/fonc.2021.778989.

69. Furie, R., Khamashta, M., Merrill, J.T., Werth, V.P., Kalunian, K., Brohawn, P., Illei, G.G., Drappa, J., Wang, L., Yoo, S., and Investigators, C.D.S. (2017). Anifrolumab, an Anti-Interferon-alpha Receptor Monoclonal Antibody, in Moderate-to-Severe Systemic Lupus Erythematosus. Arthritis Rheumatol 69, 376–386. 10.1002/art.39962.

70. Morand, E.F., Furie, R., Tanaka, Y., Bruce, I.N., Askanase, A.D., Richez, C., Bae, S.C., Brohawn, P.Z., Pineda, L., Berglind, A., et al. (2020). Trial of Anifrolumab in Active Systemic Lupus Erythematosus. N Engl J Med 382, 211–221. 10.1056/NEJMoa1912196.

71. LaFleur, D.W., Nardelli, B., Tsareva, T., Mather, D., Feng, P., Semenuk, M., Taylor, K., Buergin, M., Chinchilla, D., Roshke, V., et al. (2001). Interferon-kappa, a novel type I interferon expressed in human keratinocytes. J Biol Chem 276, 39765–39771. 10.1074/jbc.M102502200.

72. Sarkar, M.K., Hile, G.A., Tsoi, L.C., Xing, X., Liu, J., Liang, Y., Berthier, C.C., Swindell, W.R., Patrick, M.T., Shao, S., et al. (2018). Photosensitivity and type I IFN responses in cutaneous lupus are driven by epidermal-derived interferon kappa. Ann Rheum Dis 77, 1653–1664. 10.1136/annrheumdis-2018-213197.

73. Donato, R., Cannon, B.R., Sorci, G., Riuzzi, F., Hsu, K., Weber, D.J., and Geczy, C.L. (2013). Functions of S100 proteins. Curr Mol Med 13, 24–57.

74. Takeda, K., Kaisho, T., and Akira, S. (2003). Toll-like receptors. Annu Rev Immunol 21, 335–376. 10.1146/annurev.immunol.21.120601.141126.

75. Takeda, K., and Akira, S. (2003). Toll receptors and pathogen resistance. Cell Microbiol 5, 143–153. 10.1046/j.1462-5822.2003.00264.x.

76. Fernandez-Ruiz, R., and Niewold, T.B. (2022). Type I Interferons in Autoimmunity. J Invest Dermatol 142, 793–803. 10.1016/j.jid.2021.11.031.

77. Zitvogel, L., Galluzzi, L., Kepp, O., Smyth, M.J., and Kroemer, G. (2015). Type I interferons in anticancer immunity. Nat Rev Immunol 15, 405–414. 10.1038/nri3845.

78. Choi, H., Kudchodkar, S.B., Reuschel, E.L., Asija, K., Borole, P., Agarwal, S., Van Gorder, L., Reed, C.C., Gulendran, G., Ramos, S., et al. (2020). Synthetic nucleic acid antibody prophylaxis confers rapid and durable protective immunity against Zika virus challenge. Hum Vaccin Immunother 16, 907–918. 10.1080/21645515.2019.1688038.

