## Supplementary Table 1 for "Cloning and functional characterization of novel human neutralizing anti-interferon-alpha and anti-interferon-beta antibodies"

**Supplemental Table 1. Nomenclature and genes of Human IFN-α**

| Human IFN Alpha Subtype Nomenclature | | Gene |
|---|---|---|
| IFN Alpha A | IFN Alpha 2a | IFNA2 |
| IFN Alpha 2 | IFN Alpha 2b | IFNA2 |
| IFN Alpha B2 | IFN Alpha 8 | IFNA8 |
| IFN Alpha C | IFN Alpha 10 | IFNA10 |
| IFN Alpha D | IFN Alpha 1 | IFNA1 |
| IFN Alpha F | IFN Alpha 21 | IFNA21 |
| IFN Alpha G | IFN Alpha 5 | IFNA5 |
| IFN Alpha H2 | IFN Alpha 14 | IFNA14 |
| IFN Alpha I | IFN Alpha 17 | IFNA17 |
| IFN Alpha J1 | IFN Alpha 7 | IFNA7 |
| IFN Alpha K | IFN Alpha 6 | IFNA6 |
| IFN Alpha 1 | IFN Alpha D | IFNA1 |
| IFN Alpha 4a | IFN Alpha M1 | IFNA4 |
| IFN Alpha 4b | IFN Alpha 4 | IFNA4 |
| IFN Alpha WA | IFN Alpha 16 | IFNA16 |
