## Supplementary Table 2 for "Cloning and functional characterization of novel human neutralizing anti-interferon-alpha and anti-interferon-beta antibodies"

**Table S2. Demographics of participants with MS and of persons living with HIV-1**

| Patient Number | Age | Sex | Gender |
|---|---|---|---|
| MS-001 | 64 | female | female |
| MS-002 | 42 | female | female |
| MS-003 | 53 | male | male |
| MS-004 | 54 | female | female |
| MS-005 | 36 | female | female |
| MS-006 | 60 | male | male |
| MS-007 | 41 | female | female |
| MS-008 | 38 | female | female |
| MS-009 | 65 | female | female |
| MS-010 | 55 | female | female |
| MS-011 | 38 | female | female |
| PEG-002 | 43 | female | female |
| PEG-006 | 54 | female | female |
| PEG-007 | 46 | female | female |
| PEG-008 | 53 | male | male |
| PEG-009 | 51 | male | male |
| PEG-010 | 44 | male | male |
| PEG-011 | 50 | male | male |
| PEG-013 | 30 | male | male |
| PEG-014 | 43 | male | male |
| PEG-019 | 39 | female | female |
| PEG-020 | 49 | male | male |
| PEG-022 | 43 | male | male |
| PEG-023 | 34 | male | male |
| PEG-024 | 57 | male | male |
| PEG-025 | 50 | male | male |
| PEG-027 | 29 | male | male |
| PEG-029 | 28 | male | male |
| PEG-032 | 45 | male | male |
