## Supplementary figures and images for "Cloning and functional characterization of novel human neutralizing anti-interferon-alpha and anti-interferon-beta antibodies"

### Supplementary Figures

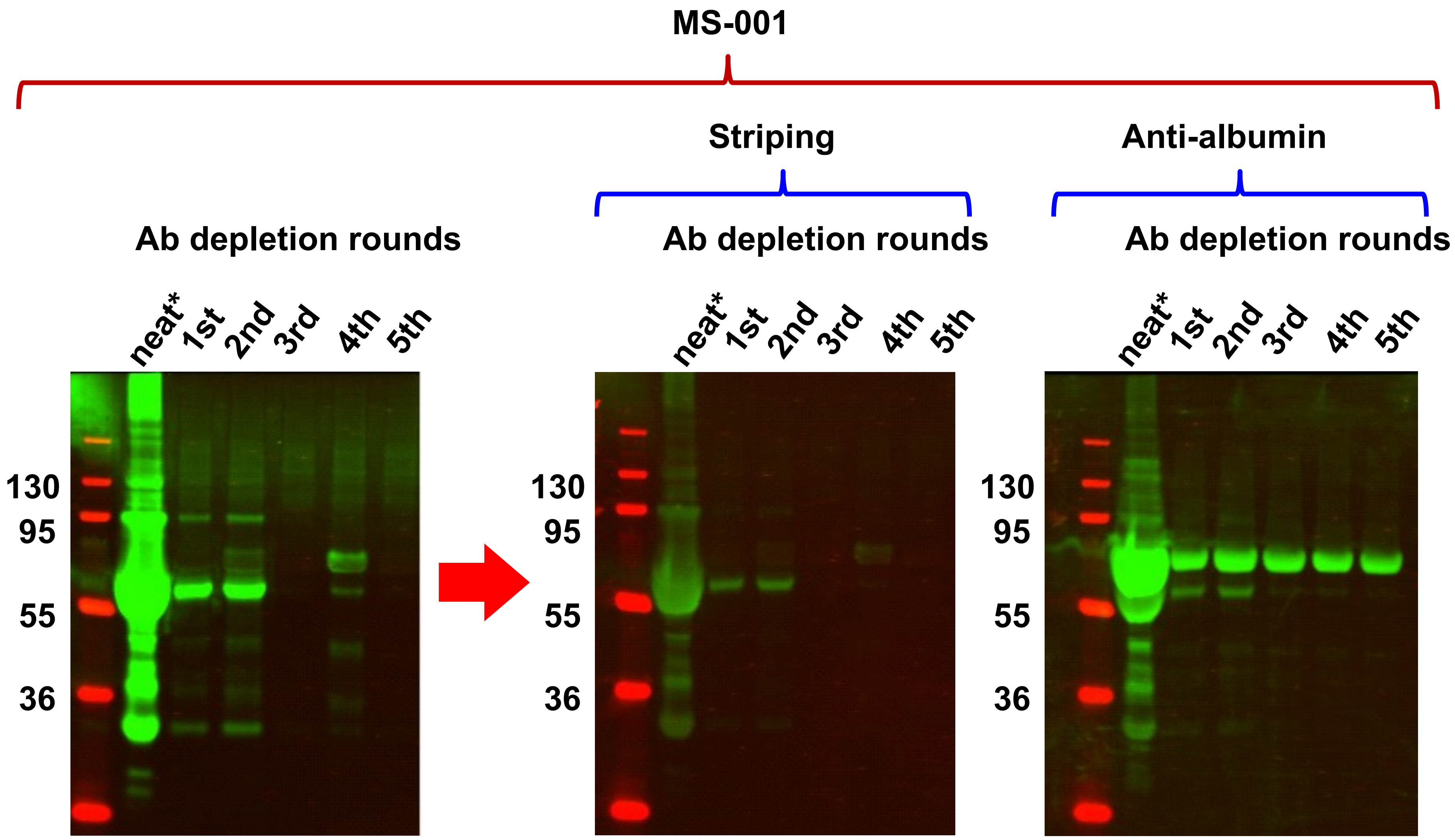

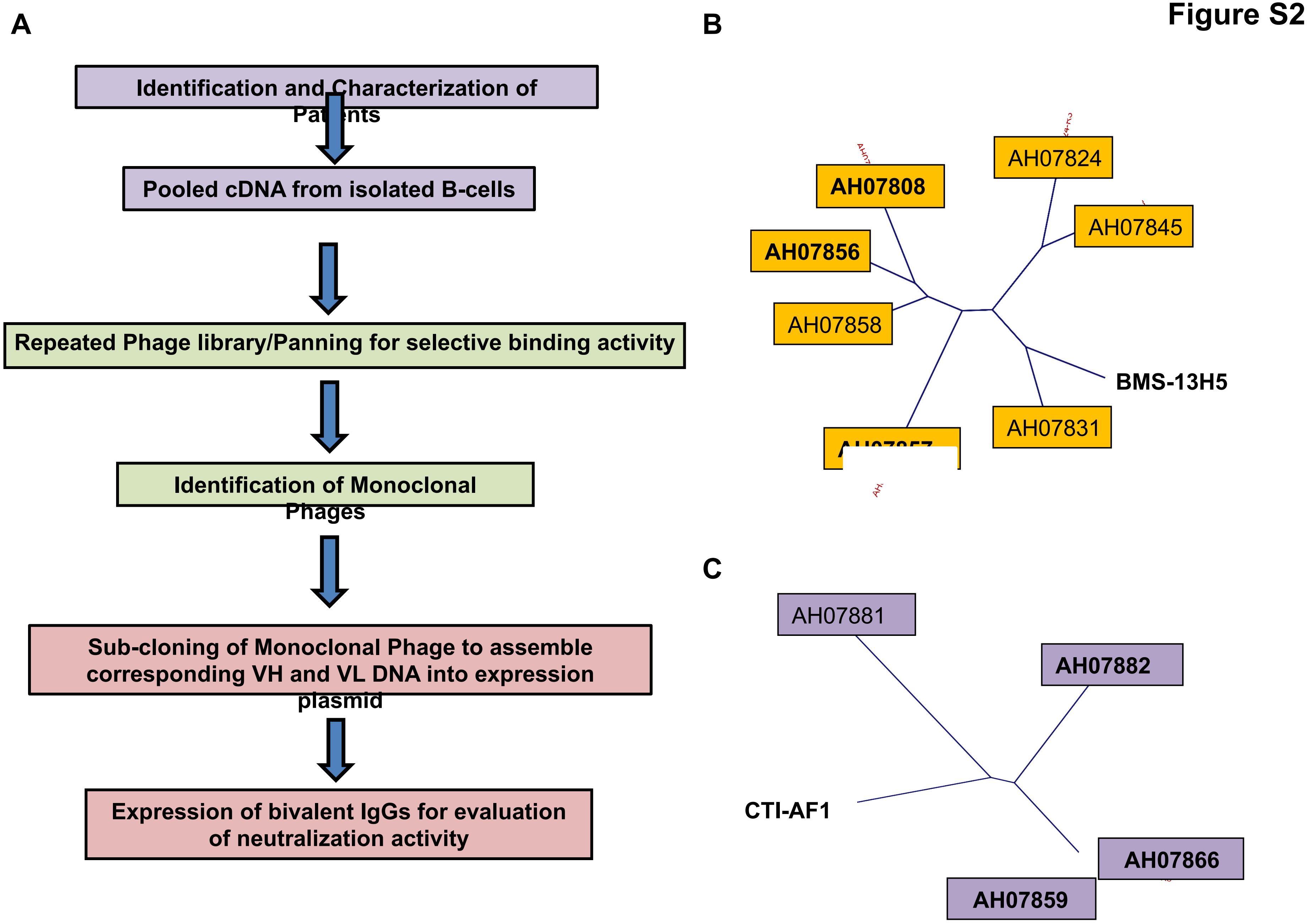

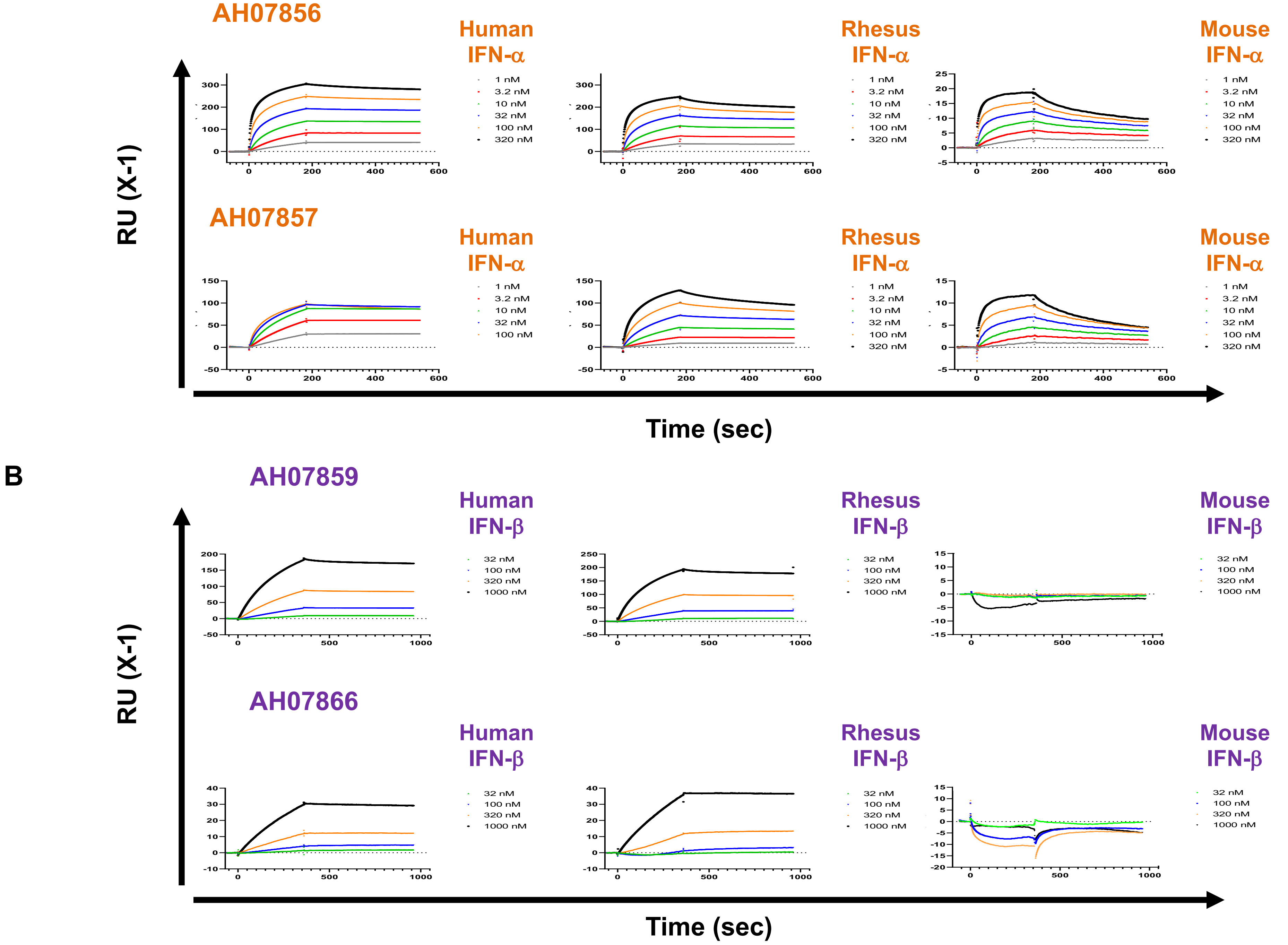

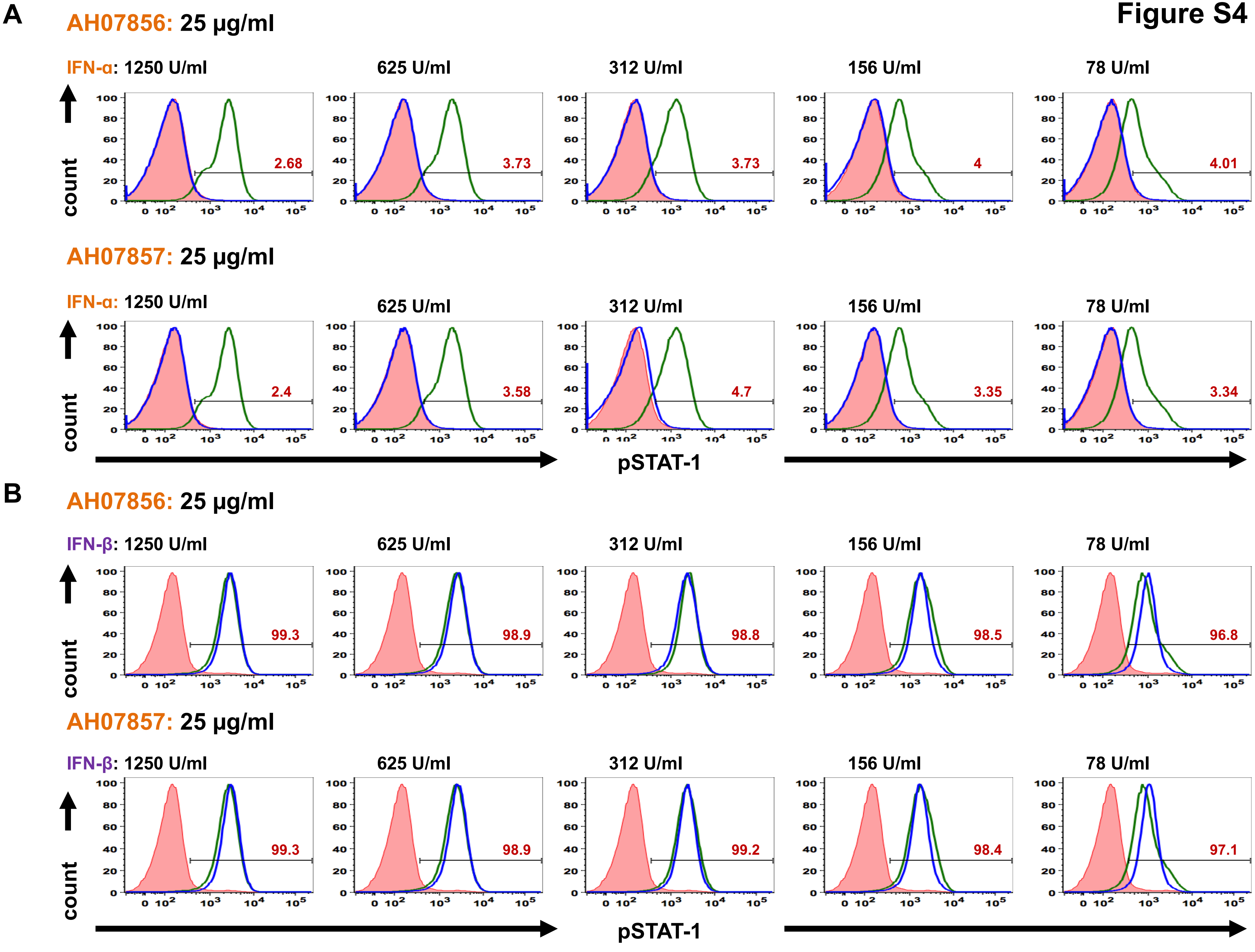

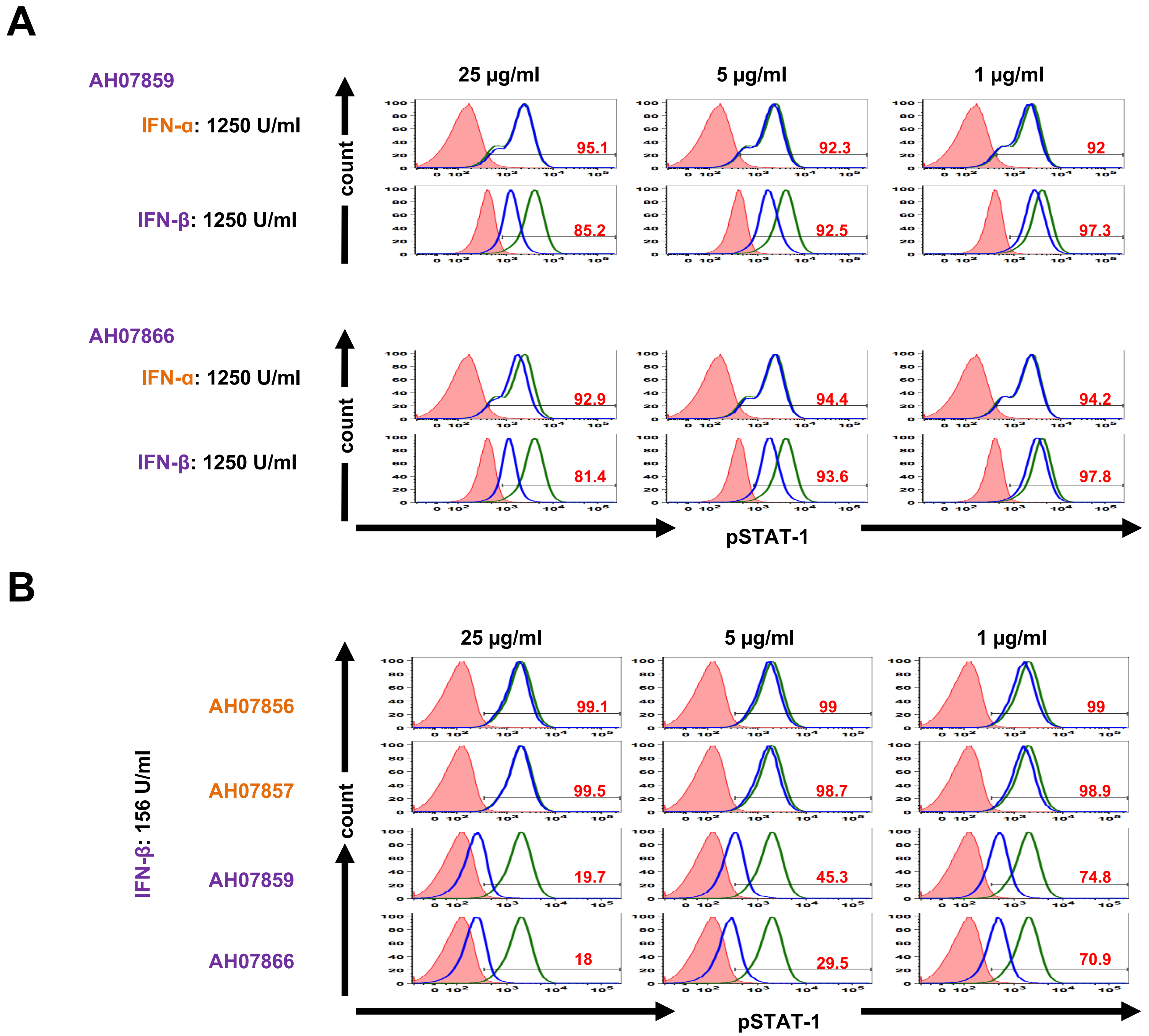

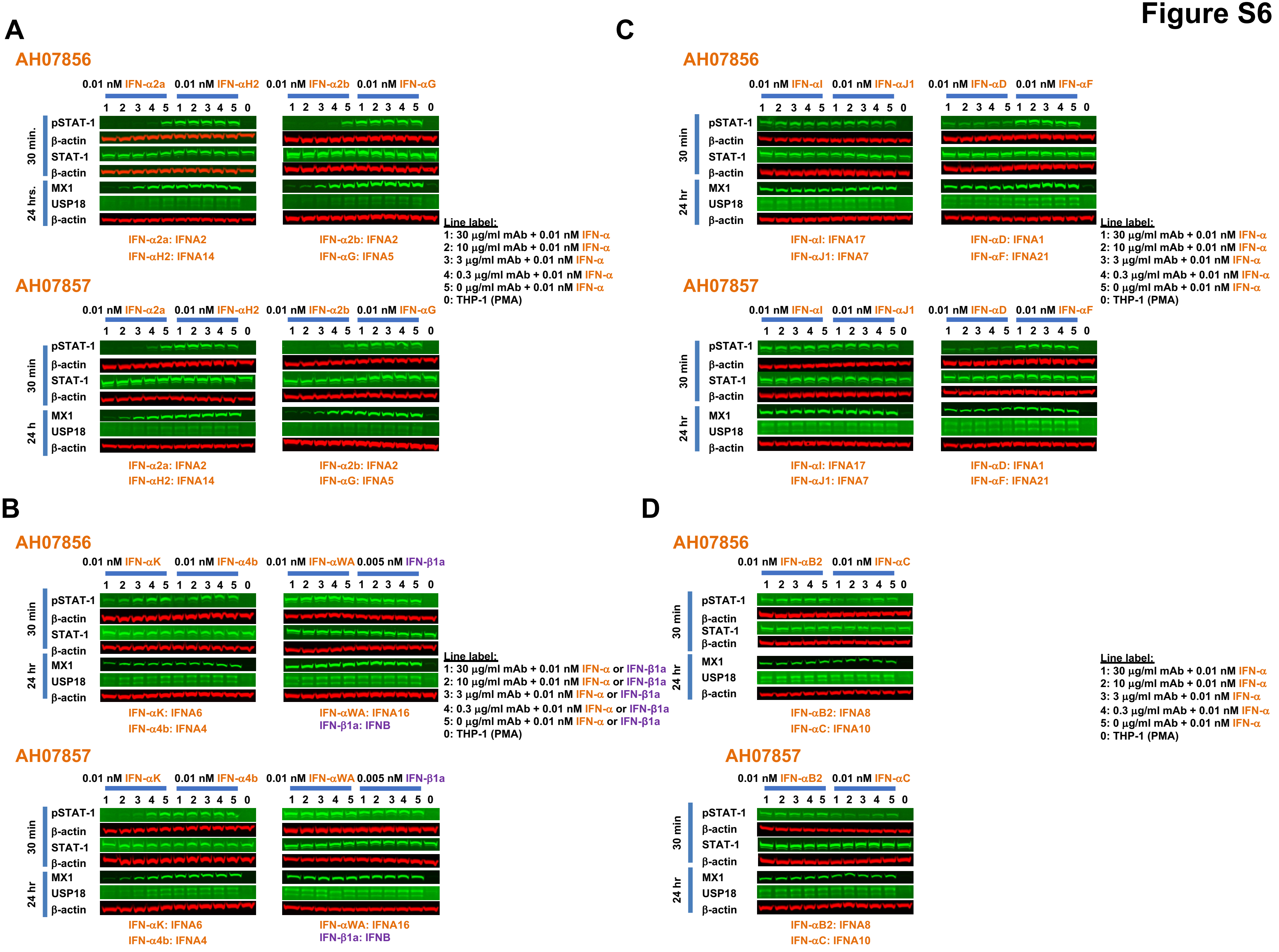

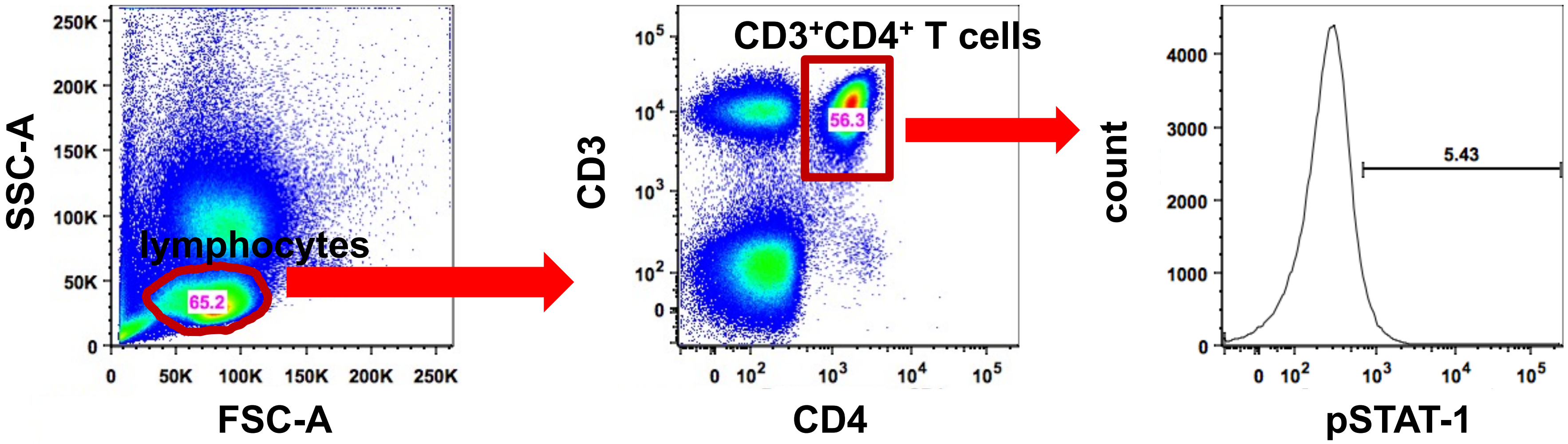
